# Alterations of Task-based fMRI Topology Underlying Cognitive Flexibility and Stability in Schizophrenia

**DOI:** 10.1101/2025.01.30.635673

**Authors:** Maren Sundermeier, Isabel Standke, Ricarda I. Schubotz, Udo Dannlowski, Rebekka Lencer, Falko Mecklenbrauck, Ima Trempler, Anoushiravan Zahedi

## Abstract

Recently, it has been suggested that brain dysconnectivity in patients with schizophrenia contributes to the wide-ranging cognitive deficits that characterize the disorder. Graph theoretical analysis offers a unique method for studying how architectural alterations in large-scale brain networks may contribute to cognitive impairments in these patients. Implementing this technique, we analyzed the functional brain activity during a predictive switch-drift task from 21 patients with schizophrenia and 22 matched healthy controls. We specifically calculated task-based global graph measures for the functional networks that were activated during expected events, events requiring a flexible updating of predictions, and events that required the stabilization of predictions. By implementing Bayesian multivariate generalized linear models, we found functional network alterations during all event types, which indicated less centralized, less integrated, and simultaneously less segregated network topology in patients with schizophrenia compared to controls. In addition, the rate of correctly detected switches, requiring flexible updating of internal models, predicted global graph measures differently for patients compared to controls. In particular, lower cognitive flexibility in patients was associated with reduced integration of functional networks. Overall, the results indicate an adaptation of network topologies, resulting in less optimal network organization in patients with schizophrenia compared to healthy controls.

## 1 Introduction

Cognitive impairments are among the core symptoms of schizophrenia (Elvevag & Goldberg, 2000). Specifically, individuals with schizophrenia suffer from poor cognitive control (Lesh et al., 2011; Orellana & Slachevsky, 2013), which is the ability to either adaptively ignore distractors (i.e., cognitive stability) or update inner representations (i.e., cognitive flexibility) based on situational demands (Dreisbach & Fröber, 2019; Trempler et al., 2017). While previous theoretical accounts and empirical investigations focused on the role of frontostriatal circuits in cognitive flexibility and stability (Arnsten & Rubia, 2012; Jiang et al., 2015; Liston et al., 2006; Miller & Cohen, 2001; Trempler et al., 2017), recent studies have shown that cognitive control is dependent on the interaction of several widespread networks (Menon & D’Esposito, 2022; Zink et al., 2021). Likewise, even though frontostriatal activity reductions that relate to cognitive control deficits have been found in schizophrenia (Cadena et al., 2018; Morey et al., 2005; Standke et al., 2021), this disorder seems better characterized by extensive brain dysconnectivity (Pettersson-Yeo et al., 2011; Zhou et al., 2015). Against this background, it seems preferable to investigate potential changes in the functional network as a whole in order to better understand cognitive deficits in patients with schizophrenia. Taking a step in this direction, in the current study, we used graph-theoretical analysis of brain imaging data to explore the topological alterations in network architecture in schizophrenia.

Graph theory, which has gained increasing popularity in the field of neuroscience (Bassett & Sporns, 2017; Bullmore & Sporns, 2009; Stam & Reijneveld, 2007), explains brain activity in terms of properties of large-scale networks, which are separated into small, densely connected communities that implement a global communication structure (Sporns, 2013). To understand these large-scale networks, one can use several different graph measures representing aspects of (I) segregation, (II) integration, (III) centrality, and (IV) resilience (Rubinov & Sporns, 2010). Measures of (I) segregation quantify the extent of functionally related regions with dense interconnections, whereas measures of (II) integration reveal how many integrative connections between segregated areas a network has, which combine specialized information. Further, (III) the centrality of a network reflects the tendency of its nodes to communicate with others. And finally, (IV) resilience assesses the network’s ability to cope with disruptions or damage.

So far, however, topological analyses of schizophrenia have yielded mixed results (Fornito et al., 2012; Kambeitz et al., 2016). The majority of studies implementing graph theory for fMRI data of these patients focus on resting-state data (e.g., Alexander-Bloch et al., 2010; Liu et al., 2008; Trempler et al., 2018; van den Heuvel et al., 2010; Yu et al., 2012; Yu et al., 2015), while only a few studies have analyzed task-based data of cognitive control or any of its components (e.g., Emmanuel D. Meram et al., 2023; Lindsay D. Oliver et al., 2021; Julia M. Sheffield et al., 2015). Notably, although several studies reported alterations of functional network topology in schizophrenia for all investigated graph measures (He et al., 2012; Ray et al., 2017; Yu et al., 2011), others only found group differences in particular measures (Ma et al., 2012; Julia M. Sheffield et al., 2015; Wang et al., 2022; Yang et al., 2020), or no effects at all (Fornito et al., 2011; Lindsay D. Oliver et al., 2021). The significant findings of group differences suggested no clear direction of alterations for functional segregation and integration (He et al., 2012; Ma et al., 2012; Ray et al., 2017; Wang et al., 2022; Yang et al., 2020; Yu et al., 2011). Additionally, centrality has not been analyzed in this context, and only one study investigated network resilience, finding higher resilience in schizophrenia (Ray et al., 2017). In regards to cognitive control components, there have been topology investigations exclusively targeting working memory (He et al., 2012; Yang et al., 2020) or cognitive flexibility performance (Wang et al., 2022), whereas cognitive stability has not yet been investigated in patients with schizophrenia. Therefore, it is unclear whether network topology alterations during task performance depend on the process currently at play.

Furthermore, for differentiating between components of cognitive control, it is beneficial to employ a task that requires both functions at different time points. Using one task where both cognitive flexibility and stability are required during different task events makes it possible to separate the effects of general task requirements (e.g., processes associated with input perception and output production) from involved cognitive functions. Our study aims to clarify potential topology alterations in functional networks associated with schizophrenia that relate to the specific processes of stabilizing and flexibly updating internal rules. We constructed graphs based on fMRI data of patients with schizophrenia and control subjects collected during the performance of a switch-drift task (Trempler et al., 2017). In the switch- drift task, digit sequences must be monitored for rule changes (*switches*), while rule-conforming noises (*drifts*) should be ignored. In contrast to previous studies, a range of global graph measures describing different characteristics of the entire network were calculated to capture neural responses relevant to cognitive flexibility and stability. We implemented Bayesian generalized linear multivariate, multilevel modeling to test the differences between patients and controls. We expected patients with schizophrenia to significantly differ from healthy controls in global graph measures considering (a) the impairments of cognitive control in schizophrenia (Orellana & Slachevsky, 2013), (b) the wide-ranging connectivity reductions in these patients (Pettersson-Yeo et al., 2011; Zhou et al., 2015), and (c) indications of altered network topology in schizophrenia related to cognitive control (He et al., 2012; Ma et al., 2012; Ray et al., 2017; Wang et al., 2022; Yang et al., 2020; Yu et al., 2011). However, with previous task-based graph theoretical findings being contradictory and inconclusive, we had non-directional hypotheses. Finally, to investigate the association between topology and behavior, we further modeled the interaction between group differences and the behavioral measures obtained from the task for the global graph measures.

## 2 Methods

### 2.1 Participants

22 patients (36.41 ± 10.28 years, seven females) diagnosed with a schizophrenia spectrum disorder participated in the study. This included 12 patients with schizophrenia and 10 patients with schizoaffective disorder. Additionally, 22 healthy controls (38.23 ± 12.26 years, nine females) were included in the present study. All patients were recruited at the Department of Mental Health of the University Hospital Münster. Diagnoses were established at consensus conferences based on the structured Clinical Interview I for DSM-IV (SCID-I) (American Psychological Association, 1994) and further available clinical data. Patients participated under regular medication, with a mean converted chlorpromazine equivalent (CPZ’; N. C. Andreasen et al., 2010) of 639.63 mg (± 427.12). Further, participants completed several questionnaires, which are not the focus of the current study. For details regarding these questionnaires, please see Table 1 in Herkströter et al.. Herkströter et al.

**Table 1.**
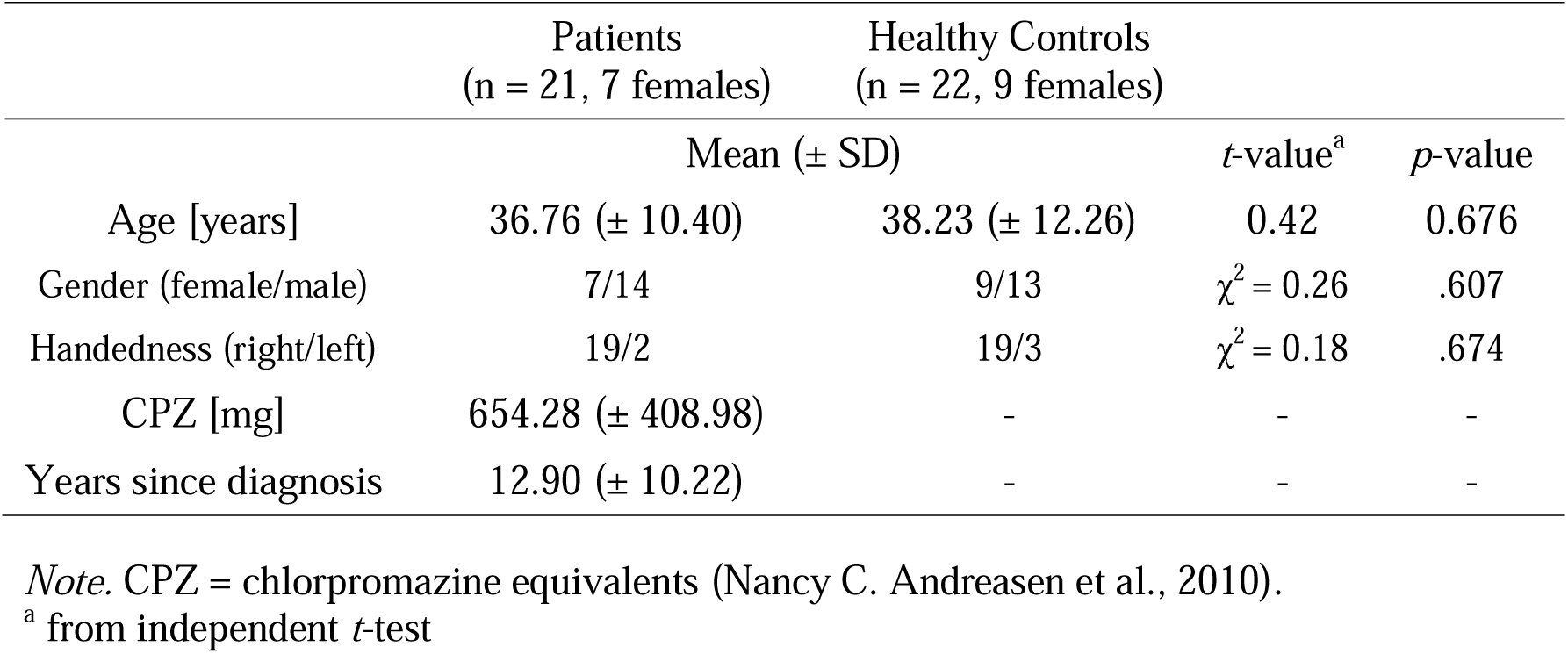
Demographical and Clinical Characteristics and Task Performance in Patients With Schizophrenia and Healthy Controls.

One patient was classified as an extreme outlier (Fig. S1) for most measures and was thus excluded from all analyses, making the final sample 21 (36.76 ± 10.40 years, seven females) patients and 22 healthy controls. Behavioral and fMRI results of the present sample were reported previously by Standke et al. (2021). Patients were recruited at the Department of Mental Health of the University Hospital Münster. Aggregated clinical data, including assessments on the Structured Clinical Interview I (SCID-I) for DSM-IV (American Psychological Association 1994), provided the basis for diagnoses confirmed at consensus conferences. Controls were checked to guarantee mental health via the short-form SCID-I screening and known history of psychotic disorders in first-degree relatives. Study procedures were approved by the local ethics committee of the University of Münster according to the Helsinki Declaration. Participants signed informed consent and were compensated for participating in the study either by monetary reimbursement or credit courses. Details on demographical and clinical characteristics are presented in Table 1.

### 2.2 Stimuli and Task

A serial predictive switch-drift paradigm (Trempler et al., 2017) was used to assess cognitive flexibility and stability. In the task, a predictable sequence consisting of four consecutive digits was presented repeatedly with a presentation time of 1 s per digit and 100 ms in between (Fig. 1). Three different types of unexpected events occurred at variable positions of the sequence during the task. Firstly, the direction of the sequence presentation could be reversed from ascending to descending or vice versa (switches). Consequently, the generated internal model had to be updated, indicating cognitive flexibility as a response to prediction errors. Participants were instructed to press a button as a reaction to this event. Secondly, omissions of single digits could appear during the sequence (drifts). Here, participants were asked not to respond, demonstrating shielding of the current internal model and, thus, cognitive stability. Lastly, motor control trials were included, where one digit was repeated up to eight times until participants reacted by pressing a button. The motor control trials provided the basis for the calculation of individual reaction time windows that were used to classify button presses as event-related or not.

**Figure 1.**
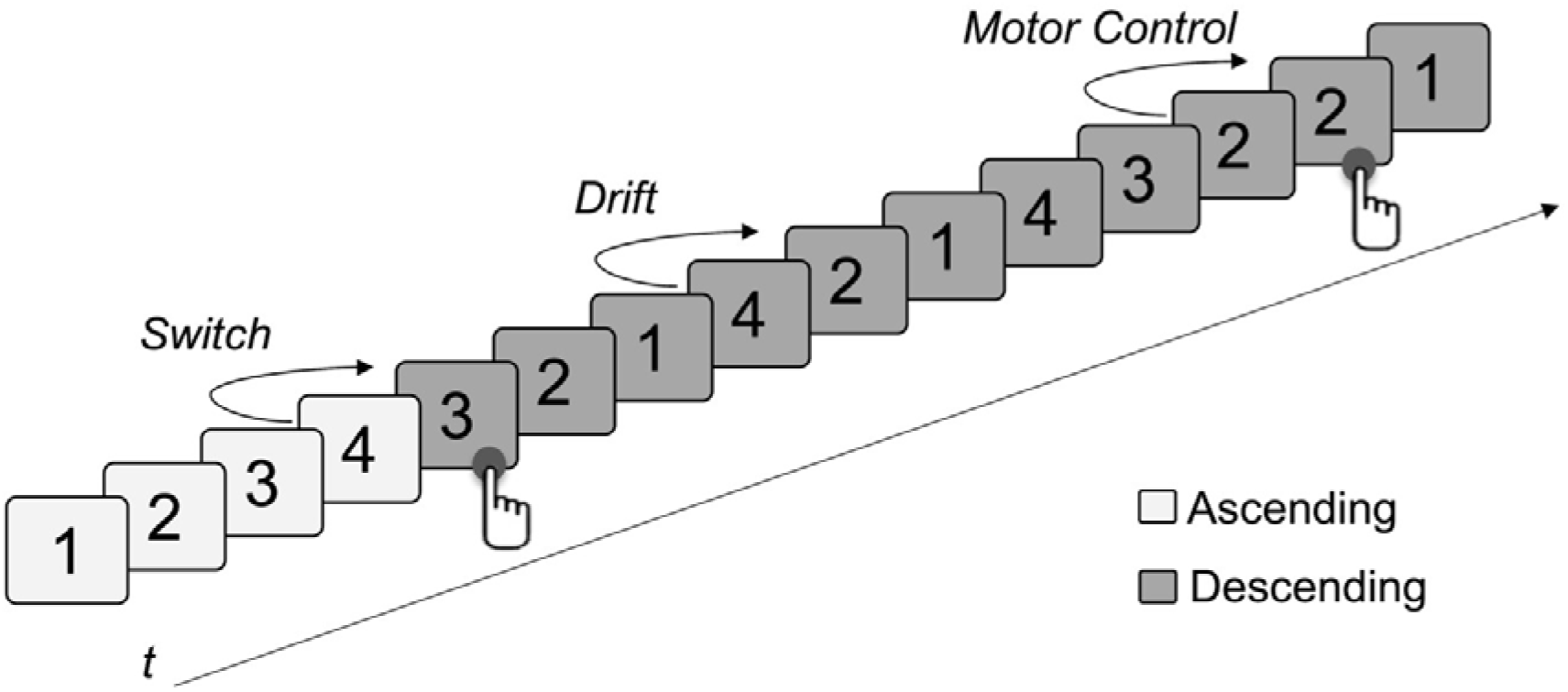
Schematic diagram of the task design. During the presentation of a predictable sequence of four consecutive digits, three different unexpected events occurred at various positions of the sequence. *Switches* represent the reversal of sequence direction and have to be indicated by a button press. *Drifts* are omissions of single digits that had to be ignored. *Motor control* trials involved the repetition of a single digit up to eight times and at least until participants reacted with a button press. These trials were used to calculate individual reaction time windows and, therefore, classify button presses during the task as event-related or not.

The task consisted of 12 blocks with an average of 125 digits per block. The different event types were distributed in a balanced way during the task using the stochastic universal sampling method (Baker, 1987). Between blocks, a fixation cross was presented for 6 s, serving as the baseline. MATLAB R2012b (The MathWorks Inc., Natick, MA, USA) was used to implement the randomization, and stimuli were presented using Presentation 18.1 (Neurobehavioral Systems, San Francisco, CA, USA). All participants completed ten blocks, each with 80 trials as practice a day before the main session. In the main session, three practice blocks were administrated directly before the task. The task was performed by the participants during fMRI data acquisition in the scanner.

### 2.3 FMRI Data Acquisition and Preprocessing

Imaging data were collected using a 20-channel head coil on a 3 Tesla Siemens Magnetom Prisma MR. Functional images were acquired using a T2*-weighted single-shot echo-planar imaging (EPI) sequence (210 mm field of view, 64×64 matrix, 90° flip angle, repetition time = 2000 ms, echo time = 30 ms). Thirty-three axial slices per volume with a slice thickness of 3 mm and a gap of 1 mm were recorded, with images being oriented parallel to the anterior commissure-posterior commissure (AC-PC) line. Structural data were collected using a standard Siemens T1-weighted magnetization-prepared rapid acquisition with gradient echo (MPRAGE) sequence for a detailed reconstruction of anatomy (field of view 256 mm, 256 x 256 matrix, 192 slices, voxel size = 1 x 1 x 1 mm³, repetition time = 2130, echo time = 2.28 ms).

Brain image preprocessing was performed using SPM12 (Wellcome Department of Imaging Neuroscience, London, UK). Images were slice-timed to the middle slice, and individual functional magnetic resonance (EPI) images were realigned to the mean image. The anatomical scan was further co-registered by rigid body transformation to the mean functional image. In order to normalize each participant’s functional scan to the Montreal Neuroimaging Institute (MNI) template brain, scans were segmented into native space tissue components. After that, spatial smoothing with a Gaussian kernel of 8 mm full width at half-maximum (FWHM) was applied. In order to remove potential confounding effects, a denoising procedure was performed on the EPI data with the CONN toolbox (Whitfield-Gabrieli & Nieto-Castanon, 2012) using the default settings. The toolbox removes confounds by implementing the anatomical component-based noise correction method (aCompCor). During denoising, the first five principal components for white matter, cerebrospinal fluid, motion parameters, and their temporal derivates were included in the model as nuisance regressors. Lastly, data was filtered with a 128 s temporal high-pass filter.

### 2.4 Graph Construction and Graph Theoretical Measures

To model event-specific functional connectivity networks, the MATLAB toolbox BASCO (Beta Series Correlation; Göttlich et al., 2015) was used to create beta series correlations (Rissman et al., 2004). This method is specifically designed to capture inter-regional interactions for event-based fMRI data (Rissman et al., 2004). Using this method, one can model functional connectivity during distinct, closely spaced stages of a cognitive task. The advantage over other approaches is that beta series correlations can tease cognitive subcomponents apart by estimating the proportion of the functional connectivity allocated to each cognitive operation. In the current study, this allows us to differentiate the functional connectivity underlying standard digit processing from functional connectivity specifically present during cognitive flexibility and stability performance. Further, beta series correlations have been used by several studies analyzing multiple components of task-based fMRI data (e.g., Fornito et al., 2011; He et al., 2012; Wang et al., 2022), which attests to the reliability of the method. The approach includes modeling the BOLD response in a general linear model (GLM) where the evoked activity in each voxel is modeled as a separate regressor for each event. Each regressor reflects brain activity using the canonical hemodynamic response function (HRF). Event length was set to zero; with the actual repetition time being 2 seconds, this allows only the immediate BOLD response to be captured. Resulting beta values can then be sorted by experimental conditions, creating corresponding beta series. Using this procedure, beta series were created for the presentation of standard digits, representing expected events, as well as switches and drifts, representing unexpected events. Only drifts and switches with a minimum distance to the next unexpected event (calculated using the participants’ specific response time windows) were included in calculating the beta series. Due to these exclusions, the number of events per participant varied slightly, but the overall number of standard digits (mean = 113.205; sd = 6.238), switches (mean = 103.886; sd= 4.363), drifts (mean = 103.682; sd = 5.552) was sufficiently large to make the variation statistically irrelevant. In other words, the differences between the number of conditions per participant were less than 10% of the total number of events per condition and participant. Afterward, the brains of all participants were parcellated according to the automated anatomical labeling (AAL) atlas (Tzourio-Mazoyer et al., 2002). Mean beta series per brain region were calculated and correlated for each pair of regions using Pearson correlation. This resulted in three correlation matrices per participant, indicating the functional connectivity between brain regions for each type of event.

Functional connectivity matrices were further processed with GraphVar (Kruschwitz et al., 2015), a MATLAB toolbox specifically designed to perform graph theoretical analyses of brain connectivity. Negative correlations were set to zero, and matrices were proportionally thresholded over a range of possible thresholds, resulting in different network sparsity levels, with the strongest links (10% to 50% with 2% increments) extracted, representing the final edges of the constructed graph. This approach is in line with recommendations from the literature (Rubinov & Sporns, 2010) and has been used by previous studies (Fornito et al., 2011; Ray et al., 2017; Yang et al., 2020). Importantly, graphs were not binarized since the inclusion of edge weights can partially compensate for differences in overall functional connectivity (van den Heuvel et al., 2017; Váša et al., 2018).

Topological measures were calculated at each threshold for the undirected, weighted connectivity matrices of each participant. This resulted in graph theoretical measures for networks that were activated during the presentation of switches, drifts, and standard digits, respectively. Global measures were calculated for each network, including *global strength* as a measure of centrality, *global characteristic path length* and *global efficiency* as measures of integration, *global clustering coefficient* and *transitivity* as measures of segregation, and *assortativity* as a measure of resilience. A comprehensive overview of the mathematical definitions of all these global graph measures can be found in Rubinov and Sporns (2010).

### 2.5 Statistical Analysis

Statistical analyses of graph measures were first run in GraphVar to check effects for all network sparsity levels. Since effects stabilized over a wide range of sparsity levels (see supplementary materials; Fig. S4 – S6), further Bayesian inference statistics were applied to graph measures calculated with the middle sparsity level of 0.3, as a reasonable sparsity level around which effects were stable. This approach allowed for a more detailed analysis using multivariate Bayesian generalized linear models. We refrained from calculating Bayesian models for all sparsity levels as the stable effects over sparsity levels indicate the benefit to be neglectable, and it is computationally not reasonably feasible. We were interested in investigating group differences as well as differences between event types and the possible interaction of these factors for all graph measures. Therefore, a Bayesian generalized linear multivariate, multilevel model was implemented with Group (i.e., Patients vs. Healthy Controls), Event Type (i.e., Standards, Drifts, and Switches), and their interaction as fixed effects of interest. The intercepts for all models were healthy controls and drift events; however, the choice of intercept does not affect the final results, as we calculated all possible contrasts using the posterior draws. Age and Gender were assumed as nuisance variables, and further, a random intercept for subjects was assumed (Model 1; Equation 1). Lastly, the residual correlation between the response variables was assumed. The residual correlation explicitly takes into account the correlations between noises of different measures due to being originating from the same source (i.e., the same participant, condition, task, etc.).

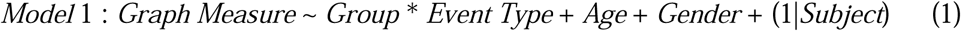

For testing the predictive capabilities of the main variables of interest, four models were compared to the full model (i.e., Model 1). The base model (Model 5, Equation 5) comprised Age and Gender as nuisance variables and assumed a random intercept for subjects. Adding to the base model, one model assumed only Group (Model 4, Equation 4), another only Event Type (Model 3, Equation 3), and the last one, both Event Type and Group as fixed effects but no interaction between them (Model 2, Equation 2).

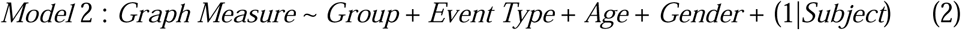

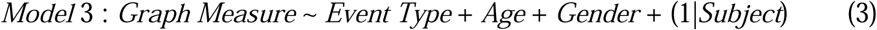

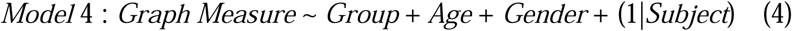

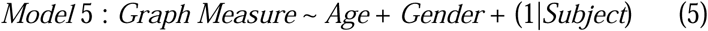

To further investigate the association between behavioral data and the graph measures, participants’ accuracy in the switch-drift task was calculated. To classify hits and false alarms, we calculated individual response time windows based on the average response times for motor control trials plus one standard deviation. Response times of under 150 ms were excluded from the analyses since those are unlikely to represent deliberate responses. Also, unexpected events, i.e., drifts and switches, that did not have a minimum distance to the next unexpected events (calculated using the participants’ specific response time window) were excluded from analyses. The rate of correctly detected switches per subject was calculated, representing the hit rate, whereas the rate of correctly ignored drifts per subject corresponds to the correct rejection rate. We fitted two Bayesian generalized linear multivariate models, including hit or correct rejection rate as a covariate of interest. One model addressed the graph measures calculated during switch processing and, therefore, included Hit Rate, Group, their interaction, as well as Age and Gender as fixed effects (Model 6, Equation 6). Equivalently, the other model addressed graph measures during drift processing and included Correct Rejection Rate, Group, their interaction, Age, and Gender as fixed effects (Model 7, Equation 7). Notably, as graph measures were calculated for different events, having a single model that included both hit rates and correct rejections is impossible, as it creates complete segregation in the data.

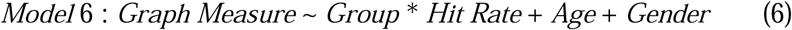

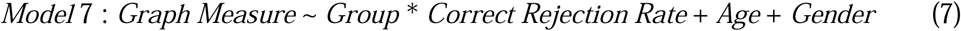

Bayesian modeling was implemented using R programming language (https://www.r-project.org/) with brms (Bürkner, 2017a) and RSTan (https://mc-stan.org/) being utilized for Bayesian modeling. We used weakly informative priors, scaling their distributions to the corresponding standard deviations of our sample, to account for the immensely varying absolute values of the different graph measures. The following priors were used for β coefficients: for global strength: N(0, 1000); for global characteristic path length: N(0, 2.5); and for global efficiency, global clustering coefficient, transitivity, and assortativity: N(0, 0.5). We used the default student-t priors for the intercepts, standard deviations, sigmas, and lkj(1) for residual correlations.

All models were fitted using four chains with 5,000 iterations each, including 2,000 iterations as warmups. If any variable showed an R-hat above 1.05 or the effective sample size was too low, the model was recalculated with increased iterations and reported accordingly. Hypotheses were tested using the hypothesis package included in brms (Bürkner, 2017b). Based on the suggestion of van Doorn et al. (2021), Bayes factors *(BF*) > 3 and BF<¾ were considered significant evidence for accepting and rejecting the tested hypothesis, respectively. All tests were two-sided (denoted by: *BF*_01_ or *BF*_10_ ) and the comparison between hypotheses and their alternative was computed via the Savage-Dickey density ratio method (Verdinelli & Wasserman, 1995). The significant Bayes factors are made **bold** for better distinguishability. For each test, we also reported the posterior probability (*p.p.*) regarding the presented hypothesis and further made it **bold** when ***p. p.*** > .95 or ***p. p***. < .05. Finally, for the model comparison, we used the Pareto smoothed importance sampling (PSIS) estimation of leave-one-out cross-validation (LOO) implemented in the loo package (Magnusson et al., 2020; Vehtari et al., 2017). LOO assesses pointwise out-of-sample prediction accuracy from a fitted Bayesian model using the log-likelihood evaluated at the posterior simulations of the parameter values; however, as it is difficult to calculate, commonly importance weights will be used instead, which results in PSIS-loo. To make sure that PSIS-loo estimation is accurate, one can use Pareto *k̑* < 0.7. However, Pareto *k̑* < 1 can still be fully trusted.

The current study uses multivariate Bayesian statistics to provide statistical analyses that are robust (Z. Dienes & McLatchie, 2018; Zoltan Dienes, 2016), and do not suffer from multiple comparison issues in the same way that a univariate frequentist analysis would do (Huberty & Morris, 1992). Therefore, the current results are reliable in terms of multiple comparisons and offer statistical insight that is independent of the sample size (Z. Dienes & McLatchie, 2018; Zoltan Dienes, 2016).

## 3 Results

### 3.1 Group and Event Effects on Global Graph Measures

Firstly, we analyzed the effects of Group, Event Type, and the interaction of these factors on the different global graph measures (Fig. 2A). For the full model (Equation 1), we used 8,000 iterations, and all chains converged with R-hats = 1.00. Further, bulk effective sample sizes were > 2,000, and tail effective sample sizes were > 1,000. Posterior predictive checks showed that the model could adequately simulate datasets with distributions representing the observed distributions for all global graph measures (Fig. 2B).

**Figure 2.**
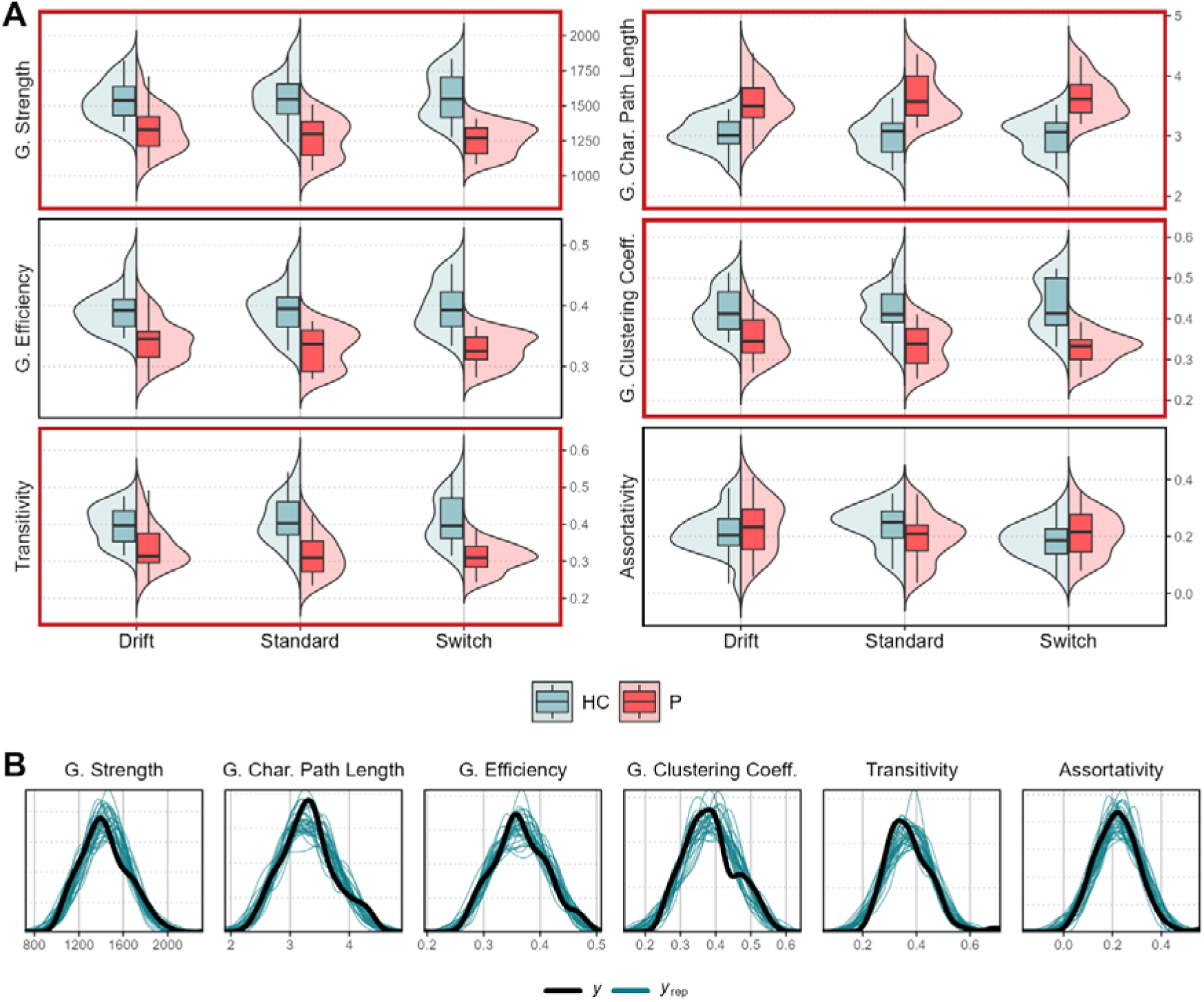
Observed and posterior predictive global (G.) graph measure distributions. **(A)** Box and violin plots of the observed global graph measure distributions for different groups and event types. Red frames indicate significant Group effects. **(B)** The density plot of the observed global graph measures distributions (*y*) and simulated distributions (*y_rep_*; Number of simulations = 100) based on the posterior predictive distributions.

To examine the effects of the variables included in the model, we tested hypotheses using the estimated posterior distributions (Fig. 3). Since we had non-directional hypotheses, all parameters were tested against the relevant null hypotheses. Table S.1 (see supplementary materials) depicts all parameter estimations and hypothesis tests for the different global graph measures. The results showed that patients with schizophrenia, compared to healthy controls, had significantly lower global strength (*H_0_*: GroupP = 0; *mean* = -217.56[-301.83, -134.74], *sd* = 42.81; ***p.p*. = 0.00**, **BF_01_ = 0.00**), higher global characteristic path length (*H_0_*: GroupP = 0; *mean* = 0.55[0.38, 0.72], *sd* = 0.09; ***p.p*. = 0.00**, **BF_01_ = 0.00**), lower global efficiency (*H_0_*: GroupP = 0; *mean* = -0.05[-0.07, -0.04], *sd* = 0.01; ***p.p*. = 0.00**, **BF_01_ = 0.00**), lower global clustering coefficient (*H_0_*: GroupP = 0; *mean* = -0.07[-0.10, -0.04], *sd* = 0.02; ***p.p*. = 0.00**, **BF_01_ = 0.00**) and lower transitivity (*H_0_*: GroupP = 0; *mean* = -0.06[-0.09, -0.03], *sd* = 0.02; *p.p*. = 0.06, **BF_01_ = 0.06**). Only in assortativity (*H_0_*: GroupP = 0; *mean* = 0.02[-0.03, 0.07], *sd* = 0.02; *p.p*. = 0.93, **BF_01_ = 14.15**), evidence suggested no group differences. Event Type did not have a main effect on graph measures and did not interact with Group either. Furthermore, post hoc contrasts using the estimated marginal means (EMMs) revealed that significant Group differences were present for all event types, i.e., patient’s graph measures for standard digits, switches, and drifts all differed significantly from those of control subjects (Table S.2).

**Figure 3.**
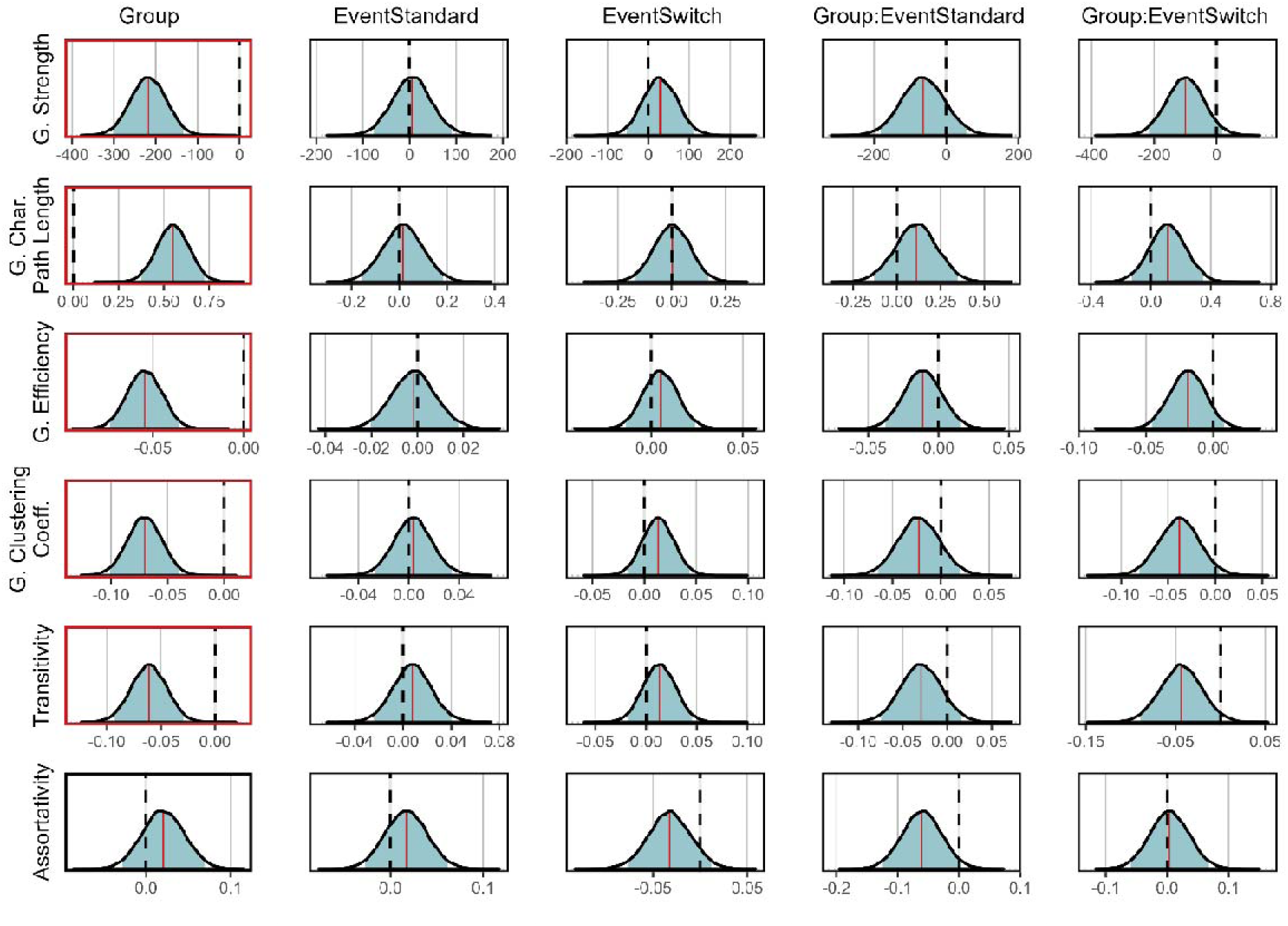
Posterior distributions of full model β coefficients. Means are indicated by the straight red lines, and 90% central posterior uncertainty intervals are indicated by blue shaded areas. The dashed black lines show null hypotheses, and red frames indicate significant effects.

As an additional step, we have investigated global graph measures based on connectivity matrices that were calculated using an alternative brain atlas that is based on a series of meta-analyses of task-related fMRI studies brain (Dosenbach et al., 2010). This was to ensure that our findings were not sensitive to the choice of brain atlas. Four participants (2 patients and 2 healthy controls) had to be excluded from this analysis since their brain imaging data did not allow for correct processing with the alternative brain atlas. Using the same multivariate Bayesian generalized linear models, we found that the results based on the Dosenbach atlas were consistent with our previous findings relying on the anatomical atlas (Fig. S3). Further, using a confirmatory analysis, we found similar results when only correct responses were included in our models instead of all responses described here (see supplementary materials for more detailed description and results).

In order to test the predictive capabilities of the fixed effects, the full model (Equation 1) was compared to four additional models (Equations 2 to 5), which systematically excluded variables. As criteria for model comparison, PSIS-loo estimations were used, which were first checked for reliability using Pareto k values of the full model. Although less than 10% of Pareto K values were above 0.7, only one was above 1, and therefore the estimations are reliable. PSIS-loo criteria clearly showed that the model only including Group (9,000 iterations used) as an effect of interest is superior against all other models (Table 2). Adding Event Type as a factor (7,000 iterations used) deteriorated the model performance by more than two standard errors. This finding is in line with the strong effect of Group that we found for the full model, as well as the absence of an effect for Event Type or their interaction.

**Table 2.** Fit Indices of the Graph Measure Models Computed Using Multivariate, Multilevel Bayesian Cumulative Modeling.

| | $\widehat{elpd}_{diff}$ | $se(\widehat{elpd}_{diff})$ | $\widehat{elpd}_{loo}$ | $se(\widehat{elpd}_{loo})$ |
| --- | --- | --- | --- | --- |
| Model 4:<br>Group + Age + Gender + (1\|Subject) | 0.0 | 0.0 | 794.6 | 26.3 |
| Model 2:<br>Group + Event Type + Age + Gender +<br>(1\|Subject) | -7.8 | 3.3 | 786.8 | 26.2 |
| Model 1:<br>Group * Event Type + Age + Gender + (1\|Subject) | -15.9 | 6.1 | 778.8 | 26.0 |
| Model 5:<br>Age + Gender + (1\|Subject) | -43.9 | 8.5 | 750.7 | 27.0 |
| Model 3:<br>Event Type + Age + Gender + (1\|Subject) | -53.6 | 8.6 | 741.0 | 26.9 |

*Note.* $\widehat{elpd}_{loo}$ = the expected log pointwise predictive density for a new dataset using the Pareto smoothed importance sampling (PSIS) leave-one-out cross-validation (LOO) criterion; $\widehat{elpd}_{diff}$ = the difference between $\widehat{elpd}_{loo}$ of two compared models; $se$ = the standard error of the targeted variable. Models are ordered by fit, with the best-fitting model at the top. All models are compared to the best-fitting model (i.e., Model 4). If a more complex model shows $\widehat{elpd}_{diff}$ greater than two and a half standard errors, it is considered to be a better fit (Magnusson et al., 2020; Vehtari et al., 2017).

### 3.2 Effects of Accuracy Data on Graph Measures

Firstly, we confirmed the previously reported significant group differences in the behavioral data (Standke et al., 2021) using Bayesian modeling (see supplementary materials for details on method) for hit rate (*H_0_*: GroupP = 0; *mean* = -0.25[-0.36, -0.15], *sd* = 0.05; *p.p*. = 0.00, **BF_01_ = 0.00**) and correct rejection rate (*H_0_*: GroupP = 0; *mean* = -0.06[-0.08, -0.04], *sd* = 0.01; *p.p*. = 0.00, **BF_01_ = 0.00**).

To investigate the association of behavioral data and the global graph measures (Fig. 4), two models were calculated, in which the accuracy from the switch-drift task was employed as a predictor (Eq. 6 & 7). For the model that included Hit Rate as a predictor for global graph measures of switch processing (Eq. 6), we found significant group differences in all global graph measures except for assortativity (Table 3). In contrast, evidence was inconclusive regarding the effect of Hit Rate for global strength (*H_0_*: Hitrate = 0; *mean* = -360.58[-693.34, -11.07], *sd* = 173.84; *p.p*. = 0.42, BF_01_ = 0.73), global characteristic path length (*H_0_*: Hitrate = 0; *mean* = 0.82[0.06, 1.57], *sd* = 0.39; *p.p*. = 0.41, BF_01_ = 0.69) and global efficiency (*H_0_*: Hitrate = 0; *mean* = -0.10[-0.18, -0.02], *sd* = 0.04; *p.p*. = 0.42, BF_01_ = 0.72) with Bayes factors below 1 but higher than ¾. However, the interaction between Group and Hit Rate was significant for global strength (*H_0_*: GroupP:Hitrate = 0; *mean* = 534.12[135.43, 925.33], *sd* = 202.46; *p.p*. = 0.14, **BF_01_ = 0.17**), global characteristic path length (*H_0_*: GroupP:Hitrate = 0; *mean* = -1.59[-2.50, -0.70], *sd* = 0.46; ***p.p*. = 0.02**, **BF_01_ = 0.02**) and global efficiency (*H_0_*: GroupP:Hitrate = 0; *mean* = 0.14[0.05, 0.23], *sd* = 0.05; *p.p*. = 0.13, **BF_01_ = 0.15**). This showed that lower values for global strength and global efficiency were associated with higher hit rates in healthy controls but with lower hit rates in patients. Similarly, a high global characteristic path length was associated with higher hit rates in control subjects but lower hit rates in patients. EMMs of linear trends revealed that global strength, global efficiency, and global characteristic path length in healthy control subjects, as well as global characteristic path length in patients, could be significantly predicted based on hit rates (Table S.3).

**Figure 4.**
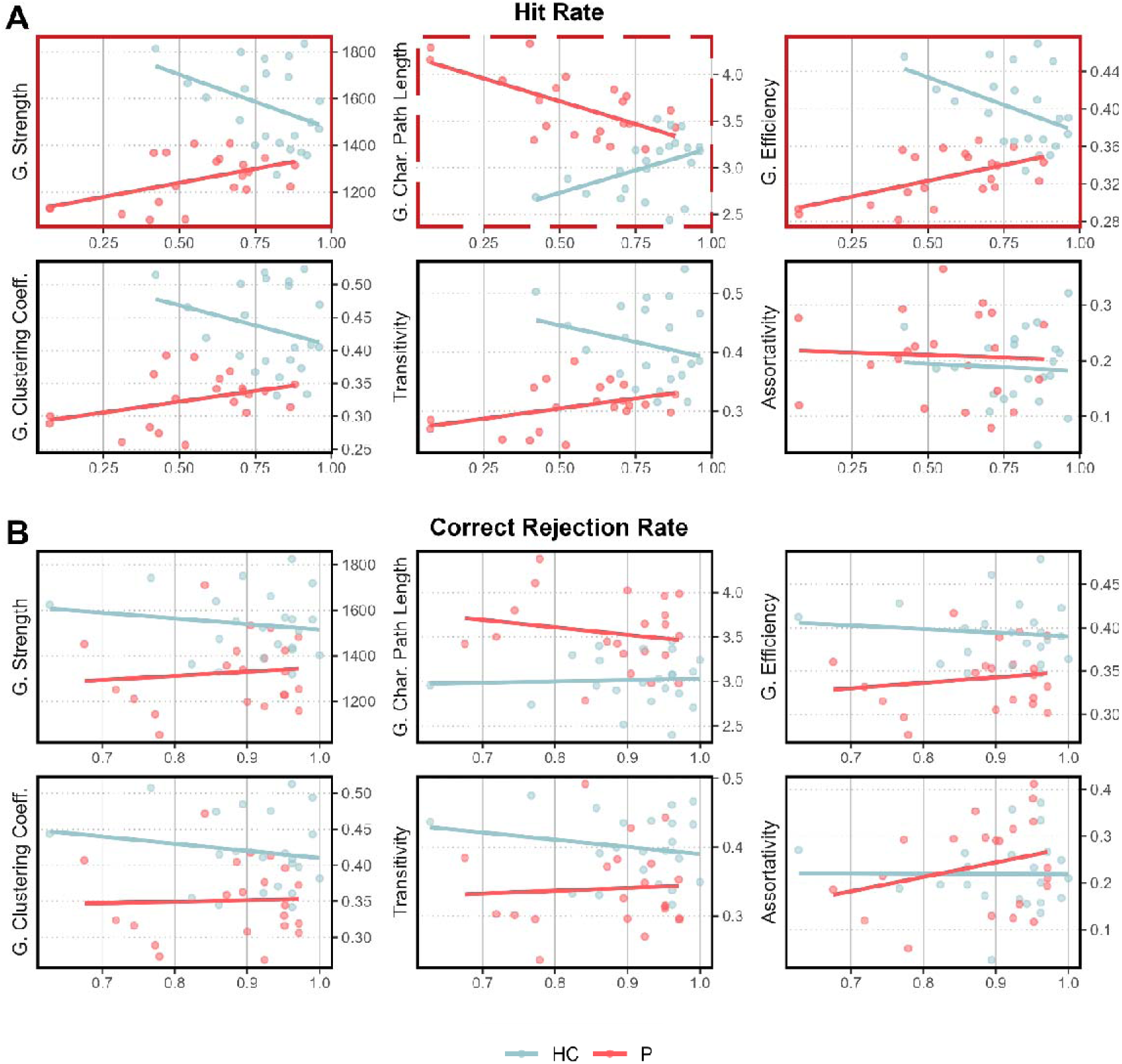
Correlations of accuracy data from the switch-drift task and global (G.) graph measures of related event types. The x-axis shows the accuracy rate, and the y-axis shows graph measure values. Red frames indicate significant slopes for healthy controls, and dashed red frames indicate significant slopes for both, healthy controls and patients. **(A)** Correlations of Hit Rate and global graph measures of switch processing for factor Group. **(B)** Correlations of Correct Rejection Rate and global graph measures of drift processing for factor Group.

**Table 3.** Results of Hit Rate as Predictor for Global Graph Measures.

|  | Coefficient | Estimate [95% CI] | E.E. | hypothesis | p.p. | B.F. |
| --- | --- | --- | --- | --- | --- | --- |
| G. Strength | GroupP | -693.98 [-976.06, -393.74] | 149.67 | GroupP = 0 | **0.00** | **0.00** |
|  | HitRate | -360.58 [-693.34, -11.07] | 173.84 | HitRate = 0 | 0.42 | 0.73 |
|  | Age | -1.29 [-4.83, 2.32] | 1.81 | Age = 0 | 1.00 | 418.21 |
|  | Gender | -1.83 [-83.72, 80.44] | 42.14 | Gender = 0 | 0.96 | 24.07 |
|  | GroupP:HitRate | 534.12 [135.43, 925.33] | 202.46 | GroupP:HitRate = 0 | 0.14 | **0.17** |
| G. Char. Path Length | GroupP | 1.73 [1.07, 2.39] | 0.34 | GroupP = 0 | **0.00** | **0.00** |
|  | HitRate | 0.82 [0.06, 1.57] | 0.39 | HitRate = 0 | 0.41 | 0.69 |
|  | Age | 0.01 [0.00, 0.01] | 0.00 | Age = 0 | 0.99 | 162.18 |
|  | Gender | 0.02 [-0.16, 0.20] | 0.09 | Gender = 0 | 0.96 | 25.80 |
|  | GroupP:HitRate | -1.59 [-2.50, -0.70] | 0.46 | GroupP:HitRate = 0 | **0.02** | **0.02** |
| G. Efficiency | GroupP | -0.17 [-0.24, -0.11] | 0.03 | GroupP = 0 | **0.00** | **0.00** |
|  | HitRate | -0.10 [-0.18, -0.02] | 0.04 | HitRate = 0 | 0.42 | 0.72 |
|  | Age | 0.00 [0.00, 0.00] | 0.00 | Age = 0 | 1.00 | 385.22 |
|  | Gender | 0.00 [-0.02, 0.02] | 0.01 | Gender = 0 | 0.98 | 51.76 |
|  | GroupP:HitRate | 0.14 [0.05, 0.23] | 0.05 | GroupP:HitRate = 0 | 0.13 | **0.15** |
| G. Clustering Coefficient | GroupP | -0.20 [-0.31, -0.08] | 0.06 | GroupP = 0 | 0.07 | **0.07** |
|  | HitRate | -0.08 [-0.21, 0.06] | 0.07 | HitRate = 0 | 0.79 | 3.68 |
|  | Age | 0.00 [0.00, 0.00] | 0.00 | Age = 0 | 1.00 | 606.99 |
|  | Gender | 0.00 [-0.04, 0.03] | 0.02 | Gender = 0 | 0.97 | 30.78 |
|  | GroupP:HitRate | 0.13 [-0.04, 0.29] | 0.08 | GroupP:HitRate = 0 | 0.65 | 1.82 |
| Transitivity | GroupP | -0.19 [-0.31, -0.07] | 0.06 | GroupP = 0 | 0.09 | **0.09** |
|  | HitRate | -0.06 [-0.20, 0.09] | 0.07 | HitRate = 0 | 0.83 | 4.90 |
|  | Age | 0.00 [0.00, 0.00] | 0.00 | Age = 0 | 1.00 | 660.24 |
|  | Gender | -0.01 [-0.04, 0.03] | 0.02 | Gender = 0 | 0.97 | 27.91 |
|  | GroupP:HitRate | 0.13 [-0.04, 0.29] | 0.08 | GroupP:HitRate = 0 | 0.65 | 1.84 |
| Assortativity | GroupP | 0.02 [-0.19, 0.23] | 0.11 | GroupP = 0 | 0.83 | 4.79 |
|  | HitRate | -0.04 [-0.28, 0.21] | 0.12 | HitRate = 0 | 0.80 | 3.99 |
|  | Age | 0.00 [0.00, 0.00] | 0.00 | Age = 0 | 1.00 | 402.21 |
|  | Gender | 0.02 [-0.04, 0.07] | 0.03 | Gender = 0 | 0.94 | 15.20 |
|  | GroupP:HitRate | -0.01 [-0.30, 0.26] | 0.14 | GroupP:HitRate = 0 | 0.78 | 3.57 |

Note. E.E. = estimation error; p.p. = posterior probability; B.F. = Bayes factor; GroupP = level Patients of factor Group. Highlighted Bayes factors and posterior probabilities indicate significant evidence for rejecting the tested hypothesis (van Doorn et al., 2021).

Further, modeling results with the model that included Correct Rejection Rate as a variable to predict graph measures during drift processing (Eq. 7) did not yield any significant effects (Table S.4).

## 4 Discussion

Aiming to gain insights into functional network topology in schizophrenia while performing a task that requires cognitive flexibility and stability, we analyzed fMRI data obtained during a switch-drift task. There is still no substantial body of literature that has investigated graph theoretical measures for understanding the neurocognitive deficits underlying schizophrenia using task-based fMRI data (e.g., Lindsay D. Oliver et al., 2021; Julia M. Sheffield et al., 2015), and the results are inconclusive. In the current study, Bayesian generalized linear multivariate, multilevel modeling revealed extensive alterations in graph measures for patients with schizophrenia compared to controls during the detection of rule changes (i.e., switches) and ignoring of rule-conforming noises (i.e., drifts). Meaningful differences were present for (I) network centrality, indicated by global strength; (II) network integration, indicated by global characteristic path length and global efficiency; and finally, (III) network segregation, measured by the global clustering coefficient and transitivity. Assortativity as a measure of network resilience was not significantly altered in schizophrenia. Analyzing the task components, there was substantial evidence that global graph measures were similar for expected digit processing, drift processing, and switch processing. Notably, we found that hit rates for correct switch detection predicted global strength, global characteristic path length, and global efficiency during switch presentation differently in patients compared to healthy subjects, while correct rejection rates did not have any predictive value for any global graph measures during drift presentation.

### 4.1 General Group Differences in Network Topology

As expected, our study revealed that patients with schizophrenia had a network topology that differed from healthy controls, not only during processing predictable events but also during processing and coping with surprising events, i.e., prediction errors. The decreased global strength that was found in patients expresses an overall lower centrality of nodes in the functional networks. Graph measures further revealed a consistent pattern of a functional network topology in patients compared to controls: patients’ network was (a) less capable of integrating information globally, as indicated by higher global characteristic path length and lower global efficiency, and simultaneously, (b) less segregated in densely interconnected specialized networks, as indicated by the lower global clustering coefficient and lower transitivity. Together, these findings indicate that functional networks in schizophrenia lack the ideal properties of a highly efficient network, i.e., a small-world network, that would promote optimal cognitive functioning (Bassett & Bullmore, 2006).

Previous studies disagreed on the direction of topological alterations associated with schizophrenia during cognitive control task completion or the existence of those altogether. While there have been investigations indicating unaltered network integration (Fornito et al., 2011; Ma et al., 2012) and segregation (Fornito et al., 2011; Ma et al., 2012; Yu et al., 2011) for cognitive control, other findings are in line with our results, revealing diminished network integration (Ray et al., 2017; Wang et al., 2022; Yu et al., 2011), and segregation (Ray et al., 2017) in patients with schizophrenia. Potential explanations for the diverging results include differences in methodology, such as the choice of the utilized task and whether graph construction was based on the whole brain or pre-selected areas.

Notably, in our study, networks in patients did not show increased resilience, as reported previously (Ray et al., 2017). Ray et al. (2017) focused their analysis on an expanded frontoparietal network, which differs from our approach since we did not pre-define a network to investigate. This could be the reason for the divergent findings in these two studies. It seems that the network active during the switch-drift task does not show group differences regarding the extent of interconnectedness among central nodes.

In contrast to former studies that focused on working memory (He et al., 2012; Yang et al., 2020) or cognitive flexibility (Wang et al., 2022) as specific components of cognitive control, our study showed that topology alterations are also present during task conditions that require cognitive stability. While we found consistent group differences for all event types, differentiating between cognitive components had no effect on topological features. Notably, our results provided substantial evidence that healthy controls similar to patients showed similar topologies when responding to different events (i.e., standard, drift, and switch events). In other words, the different event types did not cause any significant changes in topology. Corroborating our results, functional studies focusing on the loci of activation during tasks that require cognitive flexibility versus stability show that these tasks require similar areas and networks (Niendam et al., 2012; Rottschy et al., 2012). Therefore, one possible interpretation would be that differences between event types do not reveal themselves when looking at the topology of activity. In that case, one can hypothesize that the differences would be temporal rather than topological, which should be addressed in the future by a study designed to tackle temporal functional connectivity.

In summary, our results support the hypothesis that topological features in functional networks can generally be differentiated between patients and healthy controls during a task that requires both cognitive flexibility and stability. Further, these results can be interpreted as indicating that the targeted topological characteristics were relatively stable during task performance.

### 4.2 Aberrant Association of Flexibility and Global Graph Measures

Healthy controls and patients with schizophrenia differed with respect to the relationship between correct switch detection, i.e., hit rates, and global topological measures during switch processing. Specifically, we found significant group differences in the associations of hit rate with global strength, global characteristic path length, and global efficiency. When examining the effects of behavioral performance on global network measures within each group, control subjects exhibited a significant positive association between hit rate and global characteristic path length, as well as a significant negative association for global strength and global efficiency. In contrast, patients showed a significant negative association for global characteristic path length and numerically positive associations of hit rate with global strength and global efficiency. Hence, in healthy subjects, lower levels of network centrality and integration appear to facilitate cognitive flexibility performance.

Conversely, individuals with schizophrenia who exhibit better cognitive performance measured relatively higher centrality and integration levels in our data. The divergent associations of hit rate with global graph measures reflect that the biggest discrepancies in graph measures were found for the worse-performing individuals of each group. In contrast, the better-performing individuals of each group tended to have more similar global graph measures. Thus, it becomes apparent that network characteristics of centrality and integration that best supported cognitive flexibility were closer together for both groups while graph measures related to worse performance strived to diverge. This finding indicates an adaptation of network topologies in patients with schizophrenia compared to healthy controls and is of great importance for understanding the neural activity of these patients.

Previous studies in schizophrenia successfully correlated graph measures with behavioral data of cognitive functions such as working memory (Bassett et al., 2009; He et al., 2012) or an overall cognitive ability measure summarizing episodic memory, verbal memory, processing speed, goal maintenance, and visual integration (Julia M. Sheffield et al., 2017; Julia M. Sheffield et al., 2015). The reported positive associations of network integration with task performance (Bassett et al., 2009; Julia M. Sheffield et al., 2017; Julia M. Sheffield et al., 2015) agree with our results for patients with schizophrenia. Furthermore, higher network segregation was associated with lower reaction times (He et al., 2012), which is in line with our finding for patients, indicating a numerical positive relation between segregation and task performance. Less integrated and less segregated functional networks in schizophrenia, therefore, might be associated with stronger cognitive impairments over different cognitive functions. Critically, to our knowledge, group differences in associations have only been reported for resting-state fMRI data (Julia M. Sheffield et al., 2017) and not for task-based data (Julia M. Sheffield et al., 2015), while we have now shown that they can also be found in task-based data. This suggests that alterations in network topology underlying specific cognitive functions can be directly and distinguishably related to behavioral consequences. The fact that we found significant group differences in associations and a negative association between hit rate and network integration could reflect that groups greatly differ in how functional network topology supports cognitive flexibility. Our switch-drift paradigm is potentially a good candidate for differentiating associations of global graph measures and task performance between groups. However, further studies are needed in order to confirm our findings.

Interestingly, no associations and no corresponding group differences between measures of stability (i.e., correct rejection rates) and global graph measures were found. In a previous study, fMRI results from the same sample revealed reduced activity in a network comprising the inferior frontal gyrus, posterior insula, and basal ganglia when shielding against drifts (Standke et al., 2021). With these regions presumably playing a critical role in the rejection of drifts (Guo et al., 2018; Sakagami & Pan, 2007; Uddin et al., 2017), one can interpret this finding as indicating that patients did not actively reject drifts but simply missed them. This idea may explain why, in the current study, no systematic relationship was found between correct rejection rates and graph measures, and no group differences appeared. In other words, inflexibility may have contributed to the correct rejection rates in addition to stability.

### 4.3 Limitations and Future Directions

To date, there is no standard or common procedure for graph theoretical analyses, which leaves room for shortcomings in the methodology used (Hallquist & Hillary, 2019). Further studies are needed to test and confirm the influence of the different processing steps on the results. The lack of a standard procedure led to a plethora of different approaches, which might at least partly have caused the inconsistent findings in patients with schizophrenia (Kambeitz et al., 2016). Future studies would, therefore, benefit from clear guidelines similar to what is proposed for resting-state data by Hallquist and Hillary (2019) in order to make results comparable and facilitate meta-analyses.

Currently, existing graph theoretical methods cannot resolve all the challenges inherent in investigating clinical samples. As investigated by van den Heuvel et al. (2017), when applying proportional thresholds in studies including clinical samples with generally reduced functional connectivity, more spurious edges are prone to be included in the clinical sample, leading to potentially less accurate conclusions in group comparisons. In order to address this problem, we used weighted measures in the current study, but future studies should confirm our findings utilizing other approaches like probabilistic thresholding (Váša et al. (2018).

Lastly, next to our interpretation of the findings, simpler properties might have contributed to the observed effects. For instance, it cannot be ruled out that findings are at least partially based on spatial or temporal autocorrelation (Rubinov, 2023; Shinn et al., 2023). However, this does not make the differences between the two groups irrelevant, as the underlying mechanisms causing these differences can still be dissociated between the two groups. Hence, further studies are needed in order to determine what drives systematic differences in global graph measures in general, which can also provide a better interpretation of the current results.

A potential limitation of the current study is its sample size, which, while comparable to other studies (e.g., E. D. Meram et al., 2023), may be considered small. However, this concern is mitigated by the application of Bayesian statistics. By leveraging this approach, we can derive meaningful statistical insights irrespective of sample size (Z. Dienes & McLatchie, 2018; Zoltan Dienes, 2016). Notably, the use of non-informative priors (Scott & Berger, 2006) yielded robust results, as these priors allow the data to speak for itself without imposing strong assumptions. This reinforces the validity of our findings, and as such, the present results offer a significant contribution to advancing knowledge in the field.

Further, aligned with the norm in the field (Gupta et al., 2015; for review, see Haijma et al., 2013; Williams, 2008), and due to ethical and practical reasons, the patient population in the current study was not drug naïve. However, to ensure that the results of the current study were not due to drug consumption, we conducted an extra model ruling out any medication effects on the measured network characteristics (see supplementary materials for more details).

Finally, in the current study, we mainly focused on the whole brain analysis. First, unless a specific hypothesis about a particular region exists, current best practices in the field suggest that a whole-brain analysis is more appropriate (Lieberman & Cunningham, 2009; Woo et al., 2014). Moreover, network properties of the brain, by definition, cannot be fully captured within a specific ROI, as networks are inherently distributed across multiple regions (Vanhaudenhuyse et al., 2010). Given the mixed findings in previous studies (E. D. Meram et al., 2023; L. D. Oliver et al., 2021; J. M. Sheffield et al., 2015), we also intentionally refrained from focusing on a specific network, as we did not have a prior hypothesis supporting such an approach. Focusing on a specific network, however, is a valuable approach that needs to be the focus of future studies with a priori hypotheses regarding different networks.

### 4.4 Conclusion

The current study sheds light on the relationship between functional network topology and impaired cognitive flexibility and stability in schizophrenia. Our results indicate that patients with schizophrenia spectrum disorders show a less optimally organized functional network architecture characterized by reduced centrality, integration, and segregation during task performance, regardless of task event type. Hence, patients had fewer specialized local networks, and these networks showed poorer global communication and integration of information. In addition, lower network integration was associated with worse cognitive flexibility in patients. Lastly, compared to controls, patients showed opposite associations between cognitive flexibility and network centrality, integration, and segregation. These results indicate an adaptation of network topologies in patients compared to healthy controls, which deteriorated their performance regardless of whether cognitive flexibility or stability was required for the task at hand. Our findings highlight the necessity of employing a whole-brain approach to understanding cognitive deficits in schizophrenia spectrum disorders.

