## Supplementary materials for "Alterations of Task-based fMRI Topology Underlying Cognitive Flexibility and Stability in Schizophrenia"

### Supplementary Methods

#### 1. Behavioral Data Analysis

To investigate the accuracy data, Bayesian generalized linear models were implemented with Group (i.e., Patients vs. Healthy Controls) as the fixed effect of interest. Age and Gender were assumed as nuisance variables in the model for hit rate (Eq. S.1) and correct rejection rate (Eq. S.2). As in the original analysis by Standke et al. (2021), to model correct rejection rate, we further added the response bias index $B_r$ as a covariate. This response bias indicates the tendency of an individual to respond to a stimulus. It was calculated as $B_r = FA/(1-P_r)$ where $FA$ is the number of false alarms (responses to drifts) and $P_r$ is a discrimination index calculated as $P_r = H - FA$ with H being the number of hits (responses to switches).

$$Model\ 1 : Hit\ Rate \sim Group + Age + Gender \qquad \text{(S.1)}$$

$$Model\ 2 : Correct\ Rejection\ Rate \sim Group + Age + Gender + B_r \qquad \text{(S.2)}$$

#### 2. Analysis For Investigating The Effects Of Medication On Graph Measures

To investigate the effects of medication on network characteristics, a Bayesian generalized linear multivariate, multilevel model was implemented for patients only with Event Type (i.e., Standards, Drifts, and Switches) as fixed effects of interest. Notably, it is not possible to use CPZ data in a model that includes both healthy controls and patients, as CPZ data separates two groups perfectly (i.e., $CPZ > 0$ for all patients except one, and $CPZ = 0$ for healthy controls), which causes complete collinearity with the factor Group, making the model's predictions unreliable.

$$Model\ 1 : Graph\ Measure \sim Event\ Type + Age + Gender + (1|Subject) \qquad (1)$$

The results of the hypotheses checking showed that there is enough evidence to rule out the effects of medication on assortativity ($H_0: CPZ = 0; mean = 0.01[-0.02,\ 0.04], sd = 0.01; p.p. = 0.97, BF_{01} = 30.36$), clustering ($H_0: CPZ = 0; mean = -0.01[-0.02,\ 0], sd = 0.001; p.p. = 0.89, BF_{01} = 8.36$), and transitivity ($H_0: CPZ = 0; mean = -0.01[-0.02, 0.01], sd = 0.01; p.p. = 0.98, BF_{01} = 49.43$). For strength ($H_0: CPZ = 0; mean = -32.56[-56.88, -8.45], sd = 12.38; p.p. = 0.7, BF_{01} = 2.35$), efficiency ($H_0: CPZ = 0; mean = -0.01[-0.01, 0], sd = 0.01; p.p. = 0.69, BF_{01} = 2.21$), and char path ($H_0: CPZ = 0; mean = 0.08[0.02, 0.14], sd = 0.03; p.p. = 0.68, BF_{01} = 2.12$), there is only anecdotal evidence that the CPZ does not affect the network. Together, these results show that the medication cannot explain the results that have been found via the main models.

#### 3. Graph Measure Analysis for Hits and Correct Rejections

As an additional analysis, we investigated graph measures for hits and correct rejections (Fig. S7). This is to further inspect Group and Event-Type effects for trials with the same behavioral response across groups. For this analysis, five patients had to be excluded because of an insufficient number of hits (< 19), and one patient was classified as an extreme outlier later on and, therefore, excluded. Functional connectivity matrices were based on minimum if 25 hits and 25 correct rejection trials. This number was used across subject and event types to avoid confounding effects through different numbers of events. We ran the same Bayesian generalized linear multivariate, multilevel model as in our original analysis with the sole difference that this time, only correct responses were included for all the event types (Eq. 1).

$$Model\ 1 : Graph\ Measure \sim Group * Event\ Type + Age + Gender + (1|Subject) \quad (1)$$

To examine the effects of this confirmatory analysis, we used directional hypotheses where applicable based on our previous results. The results showed that patients with schizophrenia, compared to healthy controls, had significantly lower global strength ($H_0$: GroupP < 0; *mean* = -151.72[-248.86, -55.34], *sd* = 58.92; ***p.p.* = 1.00, $\mathbf{BF_{01}}$ = 192.55**), higher global characteristic path length ($H_0$: GroupP > 0; *mean* = 0.28[0.11, 0.46], *sd* = 0.11; ***p.p.* = 1.00, $\mathbf{BF_{01}}$ = 278.07**), lower global efficiency ($H_0$: GroupP < 0; *mean* = -0.04[-0.06, -0.01], *sd* = 0.01; ***p.p.* = 1.00, $\mathbf{BF_{01}}$ = 299.00**), lower global clustering coefficient ($H_0$: GroupP < 0; *mean* = -0.05[-0.08, -0.02], *sd* = 0.02; ***p.p.* = 1.00, $\mathbf{BF_{01}}$ = 239.00**) and lower transitivity ($H_0$: GroupP = 0; *mean* = -0.05[-0.09, -0.01], *sd* = 0.02; ***p.p.* = 0.99, $\mathbf{BF_{01}}$ = 79.00**). In assortativity ($H_0$: GroupP = 0; *mean* = 0.00[-0.06, 0.07], *sd* = 0.03; *p.p.* = 0.94, $BF_{01}$ = 14.70), evidence suggested no group differences. Event Type did not have a main effect on graph measures and did not interact with Group either. All these results are in line with our main analysis.

**Table S.1**

*Results of Global Graph Measure Modeling*

| | Coefficient | *Estimate [95% CI]* | *E.E.* | *hypothesis* | *p.p.* | *B.F.* |
|---|---|---|---|---|---|---|
| G. Strength | GroupP | -217.56 [-301.83, -134.74] | 42.81 | GroupP = 0 | **0.00** | **0.00** |
| | EventStandard | 5.22 [-77.73, 90.90] | 42.56 | EventStandard = 0 | 0.96 | 23.52 |
| | EventSwitch | 28.96 [-53.63, 112.57] | 42.38 | EventSwitch = 0 | 0.95 | 18.55 |
| | Age | -3.55 [-5.69, -1.36] | 1.11 | Age = 0 | 0.89 | 7.70 |
| | Gender | 16.20 [-33.52, 65.65] | 25.44 | Gender = 0 | 0.97 | 32.32 |
| | GroupP:Event Standard | -63.41 [-181.70, 56.23] | 60.40 | GroupP:Event Standard = 0 | 0.90 | 9.49 |
| | GroupP:Event Switch | -100.19 [-217.90, 19.35] | 60.37 | GroupP:Event Switch = 0 | 0.80 | 4.09 |
| G. Characteristic Path Length | GroupP | 0.55 [0.38, 0.72] | 0.09 | GroupP = 0 | **0.00** | **0.00** |
| | EventStandard | 0.02 [-0.15, 0.18] | 0.09 | EventStandard = 0 | 0.97 | 29.19 |
| | EventSwitch | 0.00 [-0.17, 0.17] | 0.09 | EventSwitch = 0 | 0.97 | 29.85 |
| | Age | 0.01 [0.01, 0.02] | 0.00 | Age = 0 | **0.00** | **0.00** |
| | Gender | -0.10 [-0.22, 0.01] | 0.06 | Gender = 0 | 0.90 | 8.86 |
| | GroupP:Event Standard | 0.11 [-0.13, 0.35] | 0.12 | GroupP:Event Standard = 0 | 0.93 | 13.31 |
| | GroupP:Event Switch | 0.11 [-0.13, 0.35] | 0.12 | GroupP:Event Switch = 0 | 0.93 | 13.39 |
| G. Efficiency | GroupP | -0.05 [-0.07, -0.04] | 0.01 | GroupP = 0 | **0.00** | **0.00** |
| | EventStandard | 0.00 [-0.02, 0.02] | 0.01 | EventStandard = 0 | 0.98 | 53.12 |
| | EventSwitch | 0.00 [-0.01, 0.02] | 0.01 | EventSwitch = 0 | 0.98 | 46.06 |
| | Age | 0.00 [0.00, 0.00] | 0.00 | Age = 0 | **0.04** | **0.05** |
| | Gender | 0.01 [-0.01, 0.02] | 0.01 | Gender = 0 | 0.98 | 59.09 |
| | GroupP:Event Standard | -0.01 [-0.04, 0.02] | 0.01 | GroupP:Event Standard = 0 | 0.96 | 26.33 |
| | GroupP:Event Switch | -0.02 [-0.05, 0.01] | 0.01 | GroupP:Event Switch = 0 | 0.93 | 14.00 |
| G. Clustering Coefficient | GroupP | -0.07 [-0.10, -0.04] | 0.02 | GroupP = 0 | **0.00** | **0.00** |
| | EventStandard | 0.00 [-0.03, 0.03] | 0.02 | EventStandard = 0 | 0.97 | 32.36 |
| | EventSwitch | 0.01 [-0.02, 0.04] | 0.02 | EventSwitch = 0 | 0.96 | 22.34 |
| | Age | 0.00 [0.00, 0.00] | 0.00 | Age = 0 | 0.98 | 39.18 |
| | Gender | 0.00 [-0.01, 0.02] | 0.01 | Gender = 0 | 0.98 | 47.12 |
| | GroupP:Event Standard | -0.02 [-0.06, 0.02] | 0.02 | GroupP:Event Standard = 0 | 0.93 | 13.43 |
| | GroupP:Event Switch | -0.04 [-0.08, 0.00] | 0.02 | GroupP:Event Switch = 0 | 0.82 | 4.68 |
| Transitivity | GroupP | -0.06 [-0.09, -0.03] | 0.02 | GroupP = 0 | 0.06 | **0.06** |
| | EventStandard | 0.01 [-0.02, 0.04] | 0.02 | EventStandard = 0 | 0.97 | 28.10 |
| | EventSwitch | 0.01 [-0.02, 0.05] | 0.02 | EventSwitch = 0 | 0.96 | 22.26 |
| | Age | 0.00 [0.00, 0.00] | 0.00 | Age = 0 | 1.00 | 259.03 |
| | Gender | 0.00 [-0.02, 0.02] | 0.01 | Gender = 0 | 0.98 | 50.29 |
| | GroupP:Event Standard | -0.03 [-0.07, 0.02] | 0.02 | GroupP:Event Standard = 0 | 0.91 | 9.54 |

| | | | | | | |
|---|---|---|---|---|---|---|
| | GroupP:Event Switch | -0.04 [-0.09, 0.00] | 0.02 | GroupP:Event Switch = 0 | 0.78 | 3.65 |
| Assortativity | GroupP | 0.02 [-0.03, 0.07] | 0.02 | GroupP = 0 | 0.93 | 14.15 |
| | EventStandard | 0.02 [-0.03, 0.06] | 0.02 | EventStandard = 0 | 0.94 | 16.43 |
| | EventSwitch | -0.03 [-0.08, 0.01] | 0.02 | EventSwitch = 0 | 0.89 | 8.26 |
| | Age | 0.00 [0.00, 0.00] | 0.00 | Age = 0 | 1.00 | 326.20 |
| | Gender | -0.01 [-0.04, 0.03] | 0.02 | Gender = 0 | 0.96 | 26.75 |
| | GroupP:Event Standard | -0.06 [-0.12, 0.00] | 0.03 | GroupP:Event Standard = 0 | 0.73 | 2.66 |
| | GroupP:Event Switch | 0.00 [-0.06, 0.07] | 0.03 | GroupP:Event Switch = 0 | 0.94 | 15.41 |

*Note.* E.E. = estimation error; p.p. = posterior probability; B.F. = Bayes factor; GroupP = level Patients of factor Group. Highlighted Bayes factors and posterior probabilities indicate significant evidence for rejecting the tested hypothesis (van Doorn et al., 2021).

**Table S.2**

*Results of Post hoc Contrasts of Estimated Marginal Means for Group Differences in Global Graph Measures*

| | Contrast | *EMM* | *Lower HPD* | *Upper HPD* |
|---|---|---|---|---|
| G. Strength | HC Standard - P Standard | **281.17** | 196.72 | 364.36 |
| | HC Switches - P Switches | **317.49** | 234.98 | 401.75 |
| | HC Drifts - P Drifts | **217.54** | 132.68 | 299.06 |
| G. Characteristic Path Length | HC Standard - P Standard | **-0.66** | -0.84 | -0.48 |
| | HC Switches - P Switches | **-0.66** | -0.84 | -0.49 |
| | HC Drifts - P Drifts | **-0.55** | -0.72 | -0.37 |
| G. Efficiency | HC Standard - P Standard | **0.07** | 0.05 | 0.08 |
| | HC Switches - P Switches | **0.07** | 0.06 | 0.09 |
| | HC Drifts - P Drifts | **0.05** | 0.04 | 0.07 |
| G. Clustering Coefficient | HC Standard - P Standard | **0.09** | 0.06 | 0.12 |
| | HC Switches - P Switches | **0.11** | 0.08 | 0.14 |
| | HC Drifts - P Drifts | **0.07** | 0.04 | 0.10 |
| Transitivity | HC Standard - P Standard | **0.09** | 0.06 | 0.12 |
| | HC Switches - P Switches | **0.10** | 0.07 | 0.14 |
| | HC Drifts - P Drifts | **0.06** | 0.03 | 0.09 |
| Assortativity | HC Standard - P Standard | 0.04 | -0.01 | 0.10 |
| | HC Switches - P Switches | -0.02 | -0.07 | 0.03 |
| | HC Drifts - P Drifts | -0.02 | -0.07 | 0.03 |

*Note.* EMM = estimated marginal mean; HC = healthy controls; P = patients. Highlighted EMMs indicate that HPD intervals do not include zero.

**Table S.3**

*Estimated Marginal Means for Linear Trends of Hit Rate and Global Graph Measures*

| | Group | *EMM* | *Lower HPD* | *Upper HPD* |
|---|---|---|---|---|
| G. Strength | Healthy Controls | **-363.5543** | -705.9607 | -26.0639 |
| | Patients | 174.5048 | -108.7622 | 446.1904 |
| G. Characteristic Path Length | Healthy Controls | **0.8237** | 0.0516 | 1.5587 |
| | Patients | **-0.7737** | -1.3848 | -0.1651 |
| G. Efficiency | Healthy Controls | **-0.0997** | -0.1797 | -0.0215 |
| | Patients | 0.0415 | -0.0246 | 0.1033 |
| G. Clustering Coefficient | Healthy Controls | -0.0813 | -0.2119 | 0.0628 |
| | Patients | 0.0472 | -0.0667 | 0.1545 |
| Transitivity | Healthy Controls | -0.0619 | -0.2021 | 0.0816 |
| | Patients | 0.0717 | -0.0458 | 0.1824 |
| Assortativity | Healthy Controls | -0.0381 | -0.2767 | 0.2068 |
| | Patients | -0.0474 | -0.2376 | 0.1433 |

*Note.* EMM = estimated marginal mean. Highlighted EMMs indicate that HPD intervals do not include zero.

**Table S.4**

*Results of Correct Rejection Rate as Predictor for Graph Measures*

| | Coefficient | *Estimate [95% CI]* | *E.E.* | *hypothesis* | *p.p.* | *B.F.* |
|---|---|---|---|---|---|---|
| Strength | GroupP | -334.25 [-893.01, 219.48] | 281.68 | GroupP = 0 | 0.65 | 1.82 |
| | CrRate | -202.82 [-704.24, 290.04] | 251.92 | CrRate = 0 | 0.74 | 2.88 |
| | Age | -4.74 [-8.16, -1.27] | 1.79 | Age = 0 | 0.94 | 15.79 |
| | Gender | 47.21 [-29.92, 125.55] | 39.75 | Gender= 0 | 0.93 | 12.64 |
| | GroupP:CrRate | 123.82 [-491.37, 745.10] | 313.58 | GroupP:CrRate = 0 | 0.75 | 2.97 |
| Characteristic Path Length | GroupP | 0.91 [-0.45, 2.28] | 0.70 | GroupP = 0 | 0.61 | 1.56 |
| | CrRate | 0.15 [-1.04, 1.34] | 0.61 | CrRate = 0 | 0.79 | 3.86 |
| | Age | 0.01 [0.00, 0.02] | 0.00 | Age = 0 | 0.84 | 5.45 |
| | Gender | -0.18 [-0.37, 0.00] | 0.09 | Gender= 0 | 0.78 | 3.57 |
| | GroupP:CrRate | -0.40 [-1.93, 1.13] | 0.78 | GroupP:CrRate = 0 | 0.73 | 2.73 |
| Efficiency | GroupP | -0.09 [-0.23, 0.04] | 0.07 | GroupP = 0 | 0.75 | 3.01 |
| | CrRate | -0.04 [-0.16, 0.08] | 0.06 | CrRate = 0 | 0.87 | 6.71 |
| | Age | 0.00 [0.00, 0.00] | 0.00 | Age = 0 | 0.92 | 12.18 |
| | Gender | 0.01 [-0.01, 0.03] | 0.01 | Gender= 0 | 0.95 | 19.15 |
| | GroupP:CrRate | 0.04 [-0.11, 0.19] | 0.08 | GroupP:CrRate = 0 | 0.85 | 5.61 |
| Clustering Coefficient | GroupP | -0.09 [-0.30, 0.13] | 0.11 | GroupP = 0 | 0.78 | 3.47 |
| | CrRate | -0.08 [-0.27, 0.11] | 0.10 | CrRate = 0 | 0.79 | 3.72 |
| | Age | 0.00 [0.00, 0.00] | 0.00 | Age = 0 | 0.98 | 65.09 |
| | Gender | 0.01 [-0.02, 0.04] | 0.01 | Gender= 0 | 0.96 | 22.49 |
| | GroupP:CrRate | 0.01 [-0.23, 0.26] | 0.12 | GroupP:CrRate = 0 | 0.80 | 4.08 |
| Transitivity | GroupP | -0.08 [-0.31, 0.15] | 0.12 | GroupP = 0 | 0.77 | 3.31 |
| | CrRate | -0.07 [-0.28, 0.13] | 0.10 | CrRate = 0 | 0.79 | 3.76 |
| | Age | 0.00 [0.00, 0.00] | 0.00 | Age = 0 | 0.99 | 124.94 |
| | Gender | 0.01 [-0.02, 0.04] | 0.02 | Gender= 0 | 0.97 | 28.28 |
| | GroupP:CrRate | 0.02 [-0.24, 0.28] | 0.13 | GroupP:CrRate = 0 | 0.79 | 3.74 |
| Assortativity | GroupP | -0.16 [-0.59, 0.25] | 0.21 | GroupP = 0 | 0.64 | 1.80 |
| | CrRate | 0.09 [-0.27, 0.47] | 0.19 | CrRate = 0 | 0.71 | 2.41 |
| | Age | 0.00 [0.00, 0.00] | 0.00 | Age = 0 | 0.99 | 112.60 |
| | Gender | -0.03 [-0.08, 0.03] | 0.03 | Gender= 0 | 0.92 | 11.53 |
| | GroupP:CrRate | 0.22 [-0.25, 0.68] | 0.24 | GroupP:CrRate = 0 | 0.59 | 1.43 |

*Note.* E.E. = estimation error; p.p. = posterior probability; B.F. = Bayes factor; GroupP = level Patients of factor Group; CrRate = Correct Rejection Rate. Highlighted Bayes factors and posterior probabilities indicate significant evidence for rejecting the tested hypothesis (van Doorn et al., 2021).

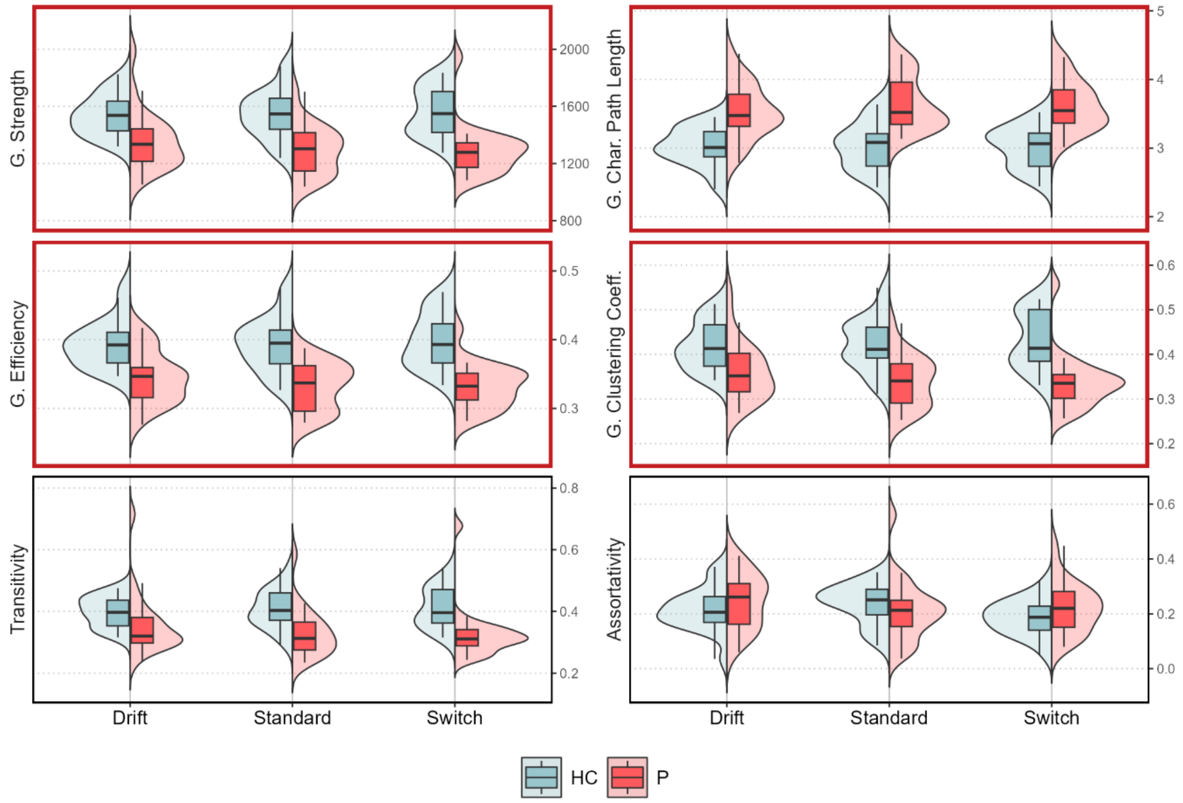


**Figure S1.** Observed global (G.) graph measure distributions for the full sample without outlier exclusion. Box and violin plots depict global graph measure distributions for different groups and event types. Red frames indicate significant Group effects.

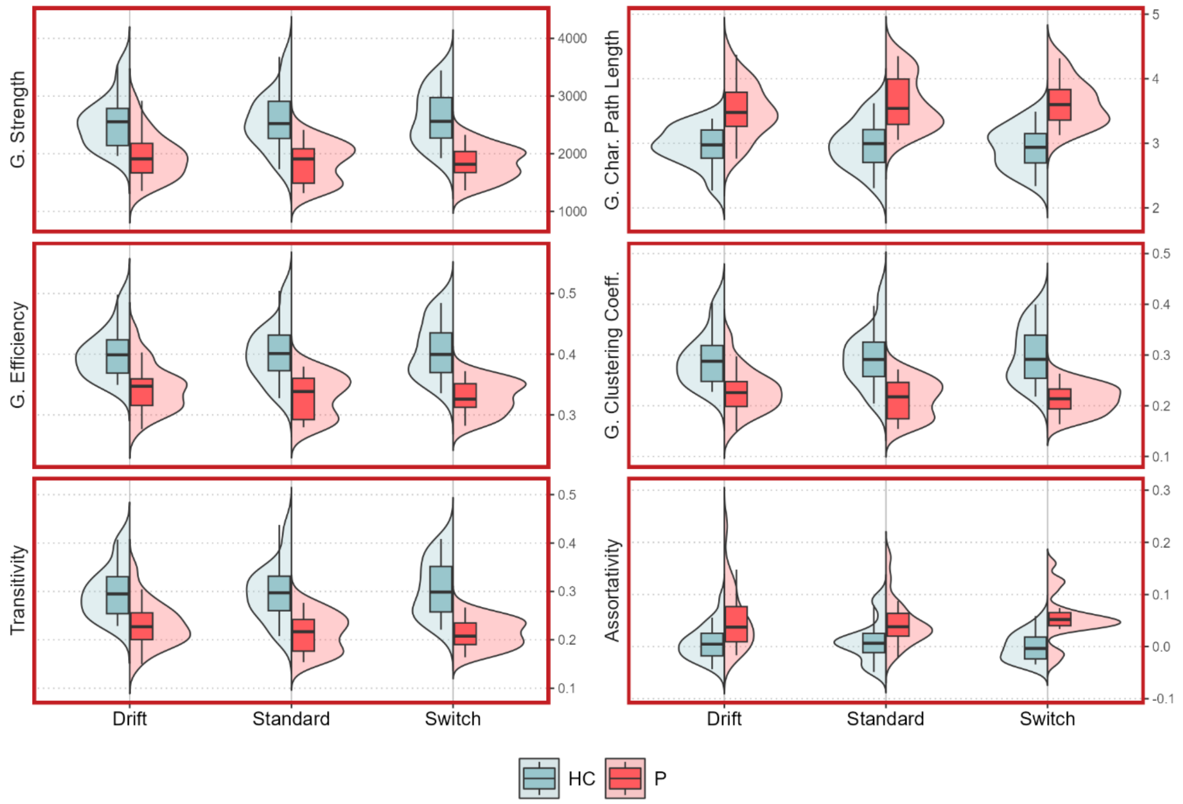


**Figure S2.** Observed global (G.) graph measure distributions for the final sample based on non-thresholded weighted connectivity matrices. Box and violin plots depict global graph measure distributions for different groups and event types. Red frames indicate significant Group effects**.**

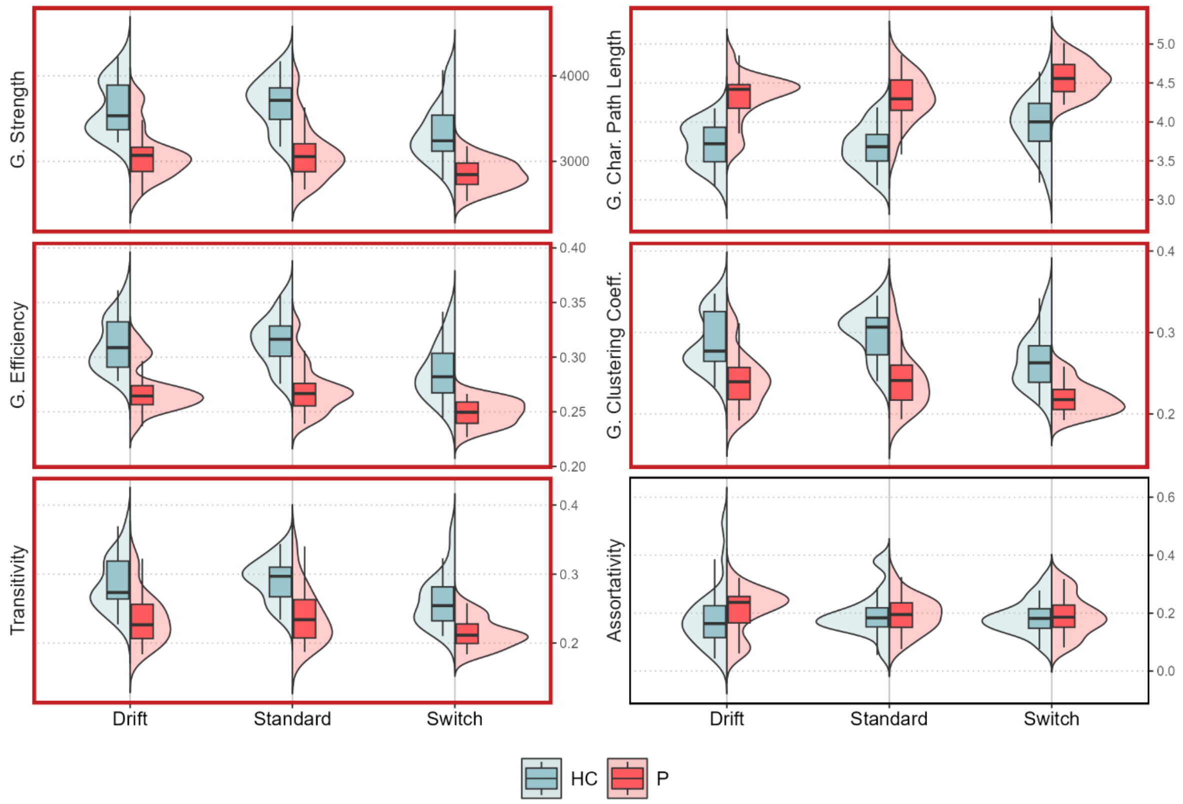


**Figure S3.** Observed global (G.) graph measure distributions for the final sample with connectivity matrices based on an alternative brain atlas (Dosenbach et al., 2010). Box and violin plots depict global graph measure distributions for different groups and event types. Red frames indicate significant Group effects**.**

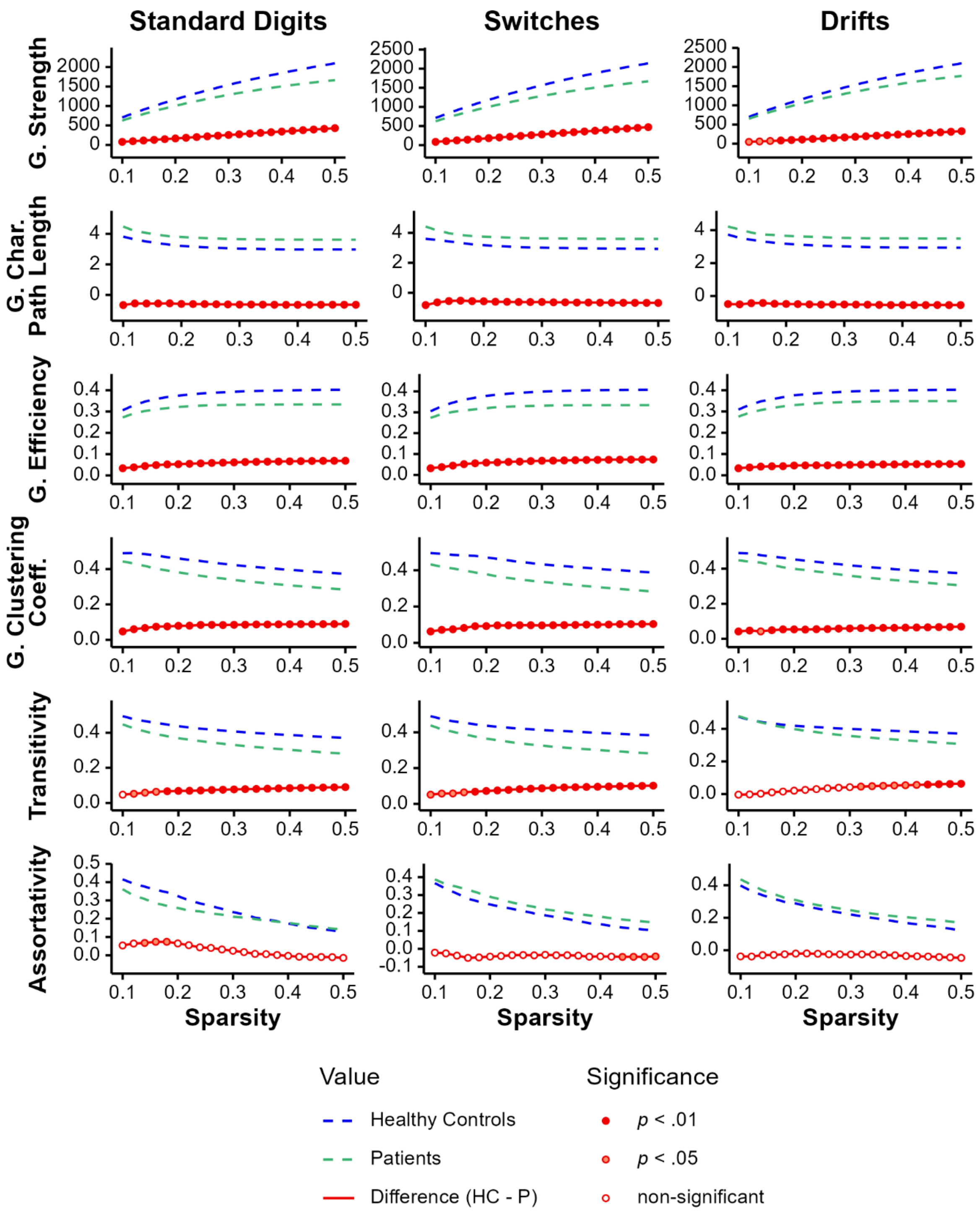


**Figure S4.** Global graph measures for different groups and event types over network sparsity levels. The difference (HC – P) describes the difference of global graph measures between healthy controls (HC) and patients (P). Significance was tested using permutation tests utilizing 10,000 repetitions.

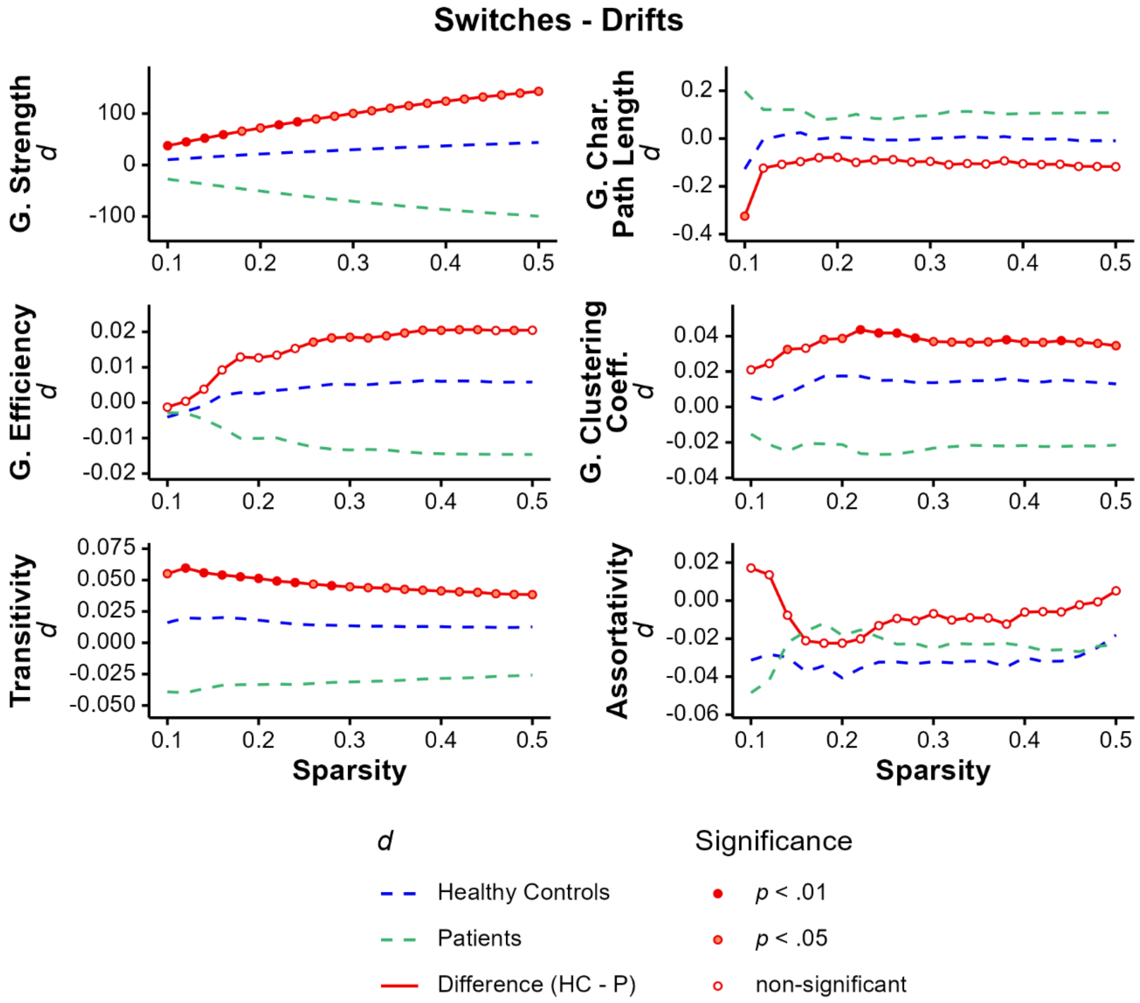


**Figure S5.** Global graph measure differences between unexpected task events for the different groups over network sparsity levels. *d* is the difference in global (G.) graph measures between switches and drifts (switches – drifts). The difference (HC – P) describes the difference of *d* between healthy controls (HC) and patients (P) and therewith the interaction of group and task events. Significance was tested using permutation tests utilizing 10,000 repetitions.

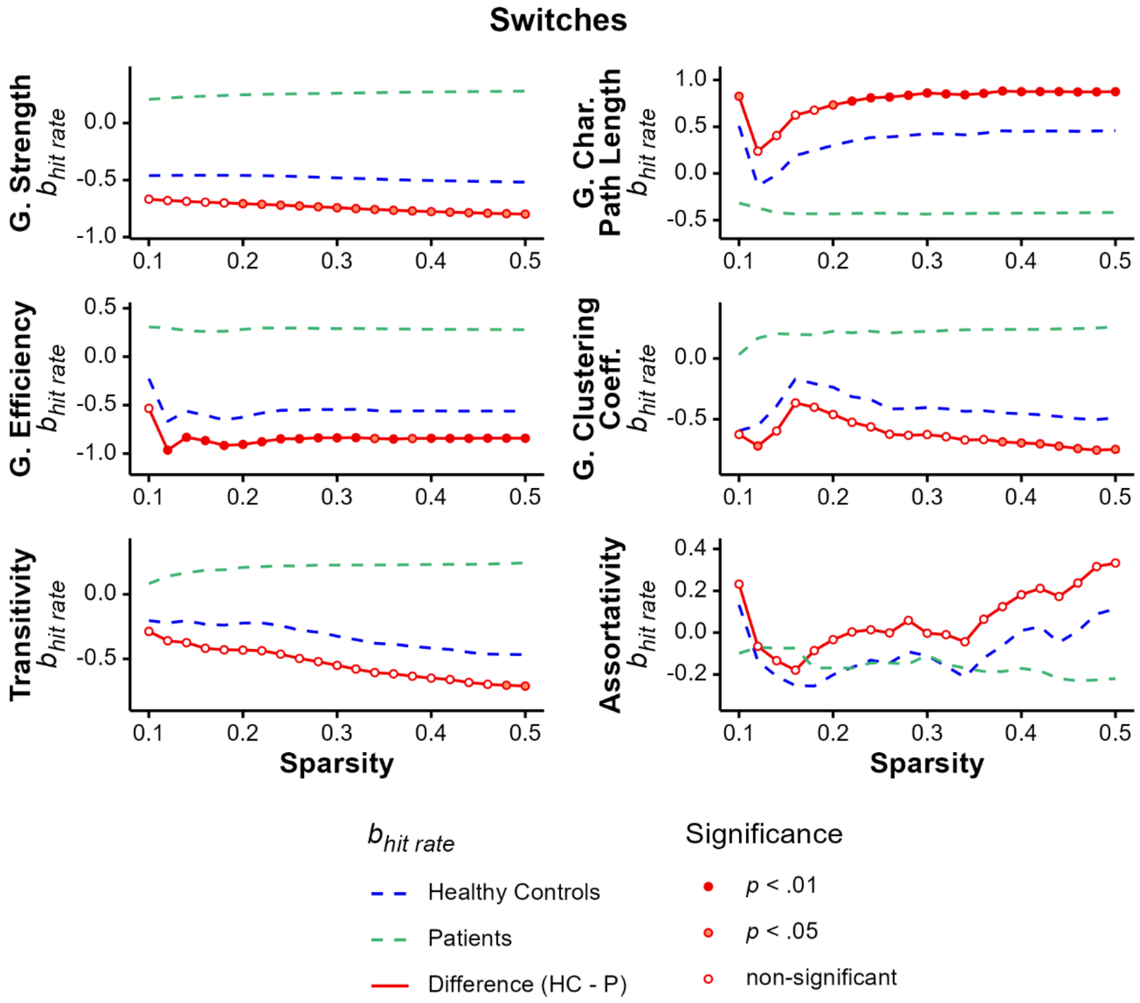


**Figure S6.** Standardized regression coefficients $b_{hit\ rate}$ of hit rate for global (G.) graph measures of switches over network sparsity levels. The difference (HC – P) describes the difference in $b_{hit\ rate}$ between healthy controls (HC) and patients (P) and therewith the interaction of group and hit rate. Significance was tested using permutation tests utilizing 10,000 repetitions.

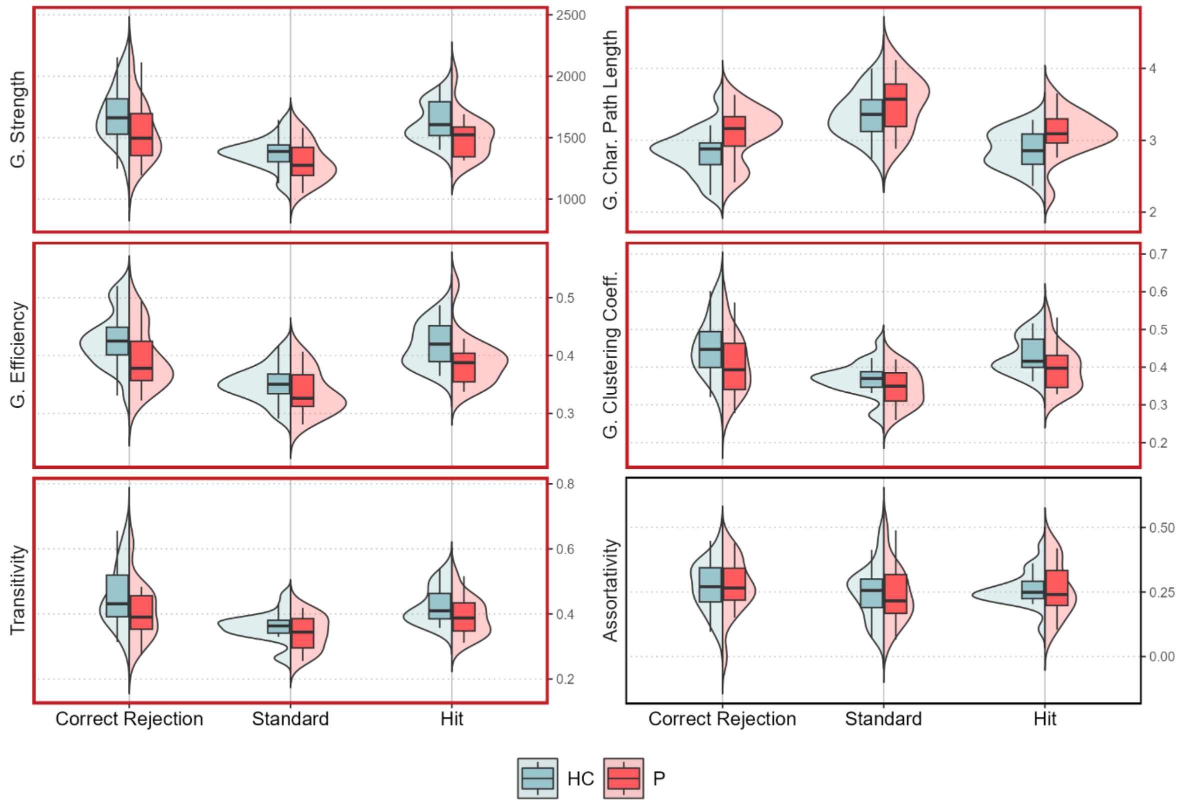


**Figure S7.** Observed global (G.) graph measure distributions for correct rejections, standard digits and hits for a subsample (17 patients 22 healthy controls). Box and violin plots depict global graph measure distributions for different groups and event types. Red frames indicate significant Group effects**.**
